# Mapping the reactive cysteine landscape of the NOD-like receptor pyrin domain-containing protein 3

**DOI:** 10.64898/2026.11.06.764924

**Authors:** Bjoern K. Ziehr, Bruno C. Amaral, David C. Schriemer, Justin A. MacDonald

**Author notes:** Correspondence to: J.A. MacDonald, Department of Biochemistry & Molecular Biology, Cumming School of Medicine, University of Calgary.

## Abstract

NOD-like receptor pyrin domain-containing protein 3 (NLRP3) is a cytosolic pattern recognition receptor that drives assembly of an oligomeric inflammasome complex to stimulate pro-inflammatory cytokine production and pyroptotic cell death. Extensive research has shown that NLRP3 activation is dependent on intracellular reactive oxygen species (ROS). Despite this, the precise targets of ROS in this process remain undefined. In this study, a combination of chemoproteomics, *in vitro* biochemical assays and molecular dynamics simulations were used to characterize reactive Cys and how they may impact NLRP3 conformation and activity status. Affinity-captured NLRP3 was subjected to reverse-thiol trapping to reveal a pool of reactive Cys residues that were identified in subsequent chemoproteomic analyses. C150, C196, and C884 were consistently detected with both *in vitro* and *in cellulo* experiments. Several NLRP3 mutants were generated; C150A and C196A (NACHT domain) as well as C933A and C941A (LRR domain) reduced IL-1β output in a reconstituted inflammasome assay. Molecular simulation of Cys sulfenylation in the LRR suggests structural alterations that reposition the trLRR loop to destabilize the inactive NLRP3 structure. Structural modelling provided insights for how Cys oxidation of the LRR may impact on NLRP3 structure. Collectively, the results establish a mechanistic understanding for how ROS may promote NLRP3 inflammasome activation.

## 1. INTRODUCTION

The innate immune system relies on genetically encoded responses to deal with diverse threats. These responses are mediated by pattern recognition receptors (PRRs) that recognize specific pathogen- and damage-associated molecular patterns (PAMP/DAMPs) [1]. The nucleotide binding and oligomerization domain (NOD)-like receptors (NLRs) are a cytoplasmic class of PRRs that have been characterized in many immunological contexts [2, 3]. NLRs containing pyrin (NLRPs) represent a subfamily of 14 members with various roles, including pro- and anti-inflammatory functions [4]. NLRP3 is the most widely studied member of this subfamily, largely due to the significant number of pathologies within which it is implicated. This includes certain cancers as well as autoimmune, gastrointestinal, cardiovascular, metabolic and neurodegenerative illnesses [5].

NLRP3 contains an N-terminal pyrin domain (PYD), a large central NACHT domain, and a C-terminal leucine rich repeat domain (LRR) [6–10]. The PYD functions as the effector domain by recruiting apoptosis associated speck-like protein containing a CARD (ASC). The NACHT domain binds ATP to drive a dramatic conformational rearrangement into the active state. Meanwhile, the LRR supports quaternary structure and participates in at least some agonist sensing. Moreover, activation of NLRP3 involves various post-translational modifications, including phosphorylation and acylation [11]. The activation and assembly of NLRP3 inflammasomes also involves numerous protein-protein interactions, several of which affect sub-cellular localization of NLRP3 [12].

NLRP3 functions to nucleate the assembly of a large multi-protein complex termed an inflammasome, which produces pro-inflammatory cytokines [13]. NLRP3 inflammasome formation begins with activated NLRP3 that forms a disk-like structure [14]. This enables recruitment of the adaptor protein ASC. Next, pro-caspase-1 is bound by ASC via CARD-CARD interactions, and the zymogen undergoes autolytic cleavage to release active heterodimers. Active caspase-1 then cleaves pro-IL-1β, pro-IL-18, and GSDMD. The N-terminal fragments of GSDMD form pores in the inner leaflet of plasma membranes which destabilizes the cell and enables secretion of IL-1β, IL-18, and other DAMPs [6, 15]. Membrane integrity continues to decline until cell lysis occurs, completing the process of pyroptosis, an inflammatory form of cell death.

Extensive research has been performed on NLRP3 to understand the regulation of this critical regulator of innate immunity. Intracellular reactive oxygen species (ROS) production is as an integral component of NLRP3 inflammasome activation [16, 17]. Generally, inhibition of ROS by chemical or genetic approaches reduces NLRP3-dependent IL-1β output in numerous cell types. However, excessive ROS and impaired reductive systems can reduce IL-1β output, highlighting the precarious nature of REDOX balance [18, 19]. Despite this complexity, ROS produced from the mitochondria and NADPH oxidases (NOX) are well established as drivers of inflammasome activation [17]. What remains unknown are the precise targets of cellular ROS that stimulate the activation of NLRP3 inflammasomes.

Cysteine residues of proteins are common and direct targets of ROS signalling pathways, and the oxidation of the Cys thiol sidechain serves as a post-translational modification (PTM) that regulates protein function [20, 21]. In its reduced form, the cysteinyl thiol is nucleophilic and displays relatively low reactivity. However, the protein microenvironment can drive thiolate (Cys-S^−^) formation at physiological pH, which is strongly nucleophilic and readily reacts with common cellular ROS such as hydrogen peroxide (H_2_O_2_), perchlorate (HOCl), and superoxide (O_2•-_) [22]. Oxidation with species like these typically results in sulfenic acid (Cys-SOH). However, other oxidized Cys species exist, including derivatives following reaction with reactive nitrogen species, termed S-nitrosothiols. Oxidized Cys-SOH residues are generally short lived, but they have been experimentally observed to directly regulate protein function [23–27].

In this study, we sought to identify ROS-sensitive Cys residues in NLRP3 and determine how those residues support inflammasome activation. Several reactive Cys were identified using *in vitro* and *in situ* chemoproteomic approaches. Additionally, *in vitro* biochemical assays and molecular dynamics simulation uncovered how certain reactive Cys may impact NLRP3 conformation and activity status. Our study establishes a novel mechanistic understanding for how NLRP3 may be directly impacted by ROS to drive inflammasome activation.

## 2. MATERIALS and METHODS

### 2.1 Materials

All chemicals were reagent grade or higher and were purchased from Fisher Scientific, Millipore-Sigma or VWR. Mouse monoclonal anti-human NLRP3 (Cat#AG-20B-0014, Adipogen, San Diego, CA), rabbit polyclonal anti-β-actin, (Cat#4967, Cell Signaling Technology, Danvers, MA), horseradish peroxidase (HRP)-conjugated goat anti-mouse IgG (Cat#115-035-003, Jackson ImmunoResearch, West Grove, PA), and donkey HRP-conjugated anti-rabbit IgG (Cat#611-7302, Rockland Immunochemicals, Limerick, PA) were used for immunoblotting. The GFP-Trap clone was a gift from Dr. Laura Trinkle-Mulcahy (University of Ottawa), and immobilized GFP-Trap Sepharose resin was produced as previously reported [28] with coupling of anti-GFP nanobody to CNBr-activated Sepharose 4B (Cytiva, Marlborough, MA). Protease Inhibitor Cocktail (cOmplete, EDTA-free), N-ethylmaleimide (NEM), 4-acetamido-4’-maleimidylstilbene-2,2’-disulfonic acid (AMS), iodoacetamide (IAM), pan-caspase inhibitor Z-VAD(OMe)-FMK (ZVAD), N-5-hexyn-1-yl-2-iodoacetamide (IAA-C6), dithiothreitol (DTT), and maleimide-PEG5K-OH (Mal-PEG5K; M_n_ 5,000) were acquired from Millipore-Sigma. The OptEIA Human IL-1β ELISA kit was from BD Bioscience (#557966). Mass spectrometry grade trypsin protease (#90058) and SuperSignal West Femto ECL substrate were purchased from ThermoFisher Scientific (Waltham, MA).

### 2.2 Molecular cloning of NLRP3-GFP Constructs

Full-length human *NLRP3* (NM_004895) in the pAcGFP-N1 vector (ClonTech/TaKaRa Bio, San Jose, Ca) was used as previously described [29]. The C8A, C108A, C8A/C108A, C150A, C514A, and C150A/C514A variants of NLRP3 were produced using a Q5 Site-directed Mutagenesis Kit (New England Biolabs, Whitby, ON) with non-overlapping primers.

### 2.3 Heterologous expression of NLRP3

HEK293T cells (ATCC# CRL-11268) were cultured in DMEM media supplemented with 10% (v/v) heat-inactivated FBS and 1% (w/v) penicillin/streptomycin. Cells were maintained on uncoated 10-cm plates in a 37 °C humidified incubator with 5% CO2. After passaging the day before transfection, cells were grown to 80% confluency and subsequently transfected with GFP-tagged NLRP3 in pAcGFP-N1 plasmid (5 μg/10 cm plate) using Lipofectamine. Media was changed after 16-18 hrs, and the transfected cells were cultured for 40 hrs before harvesting.

### 2.4 Inflammasome Reconstitution

HEK293T cells were seeded with 6.0 x 10^5^ cells/well in 24-well plates previously coated with poly-D-lysine (0.1mg/mL) The cells were grown for 24 hr prior to heterologous transfection with plasmids encoding inflammasome components: NLRP3 (40 ng), ASC (2 ng), pro-caspase-1 (20 ng), and pro-IL-1β (80 ng). After 17 h incubation, cells were gently washed with pre-warmed Dulbecco’s Phosphate-buffered Saline (DPBS). Serum-free DMEM with vehicle or treatment (10 μM nigericin) was added and cells were incubated for 1 hr. The cell culture media was collected and briefly centrifuged to remove any floating cells (1 min, 1000 x g). The resultant supernatant was used to measure IL-1β secretion by ELISA (BD Biosciences) following the manufacturer’s protocol. After discarding the remaining culture media, RIPA lysis buffer was added to the wells to harvest cellular material. Plates were frozen at −80 °C for later analyses. After thawing, cell suspensions were collected, sonicated (1x 3 s pulse), centrifuged (16,000 g, 15 min, 4 °C) and the cleared lysate was collected.

### 2.5 Reverse thiol trapping to map NLRP3 reactive cysteine content

Following ectopic expression of NLRP3-GFP, HEK293T cells were washed with phosphate-buffered saline (PBS) twice. Whole cell lysates were prepared with addition of lysis buffer (300 μL of 50 mM Tris-HCl pH 7.2, 150 mM NaCl, 0.1% NP-40 with protease inhibitor cocktail). Cleared lysate (from one 10-cm plate) was incubated with 120 μL of pre-washed GFP-Trap resin (50:50 slurry in PBS) for 1 hr at 4 °C with end-over-end agitation. Samples (10 μL of bead slurry charged with NLRP3-GFP) were oxidized with addition of CuCl_2_ (0.01 – 25 μM) for 10 min and then quenched with EDTA (200 μM). Samples were buffer-exchanged into basic buffer (pH 8.5, 50 mM Tris-HCl, 50 mM NaCl) and incubated with 20 mM IAM for 30 min. Samples were washed into pH 7.5 buffer (50 mM Tris-HCl, 50 mM NaCl) and incubated with 25 mM DTT for 15 min. Samples were then washed three times with pH 7.5 buffer and incubated with 0.5 mM Mal-PEG5K for 30 min. For AMS labelling, the experiment was performed as above, except thiols were reacted for 30 min with 1 mM AMS. All incubations were performed at 30 °C with 500 rpm shaking. NLRP3-GFP was eluted with SDS-PAGE Sample Buffer (95 °C; 10 min) and collected by brief centrifugation (30 s, 14,000x g) in a mini-spin column. Protein samples were subjected to SDS-PAGE and then anti-NLRP3 immunoblotting.

### 2.6 Western blotting

Samples were prepared for SDS-PAGE from cleared whole cell lysates following normalization of protein concentrations. Proteins were separated on 10% (w/v) SDS-PAGE gels and transferred to polyvinylidene fluoride (PVDF, 0.45 μm pore) membranes for 1 h using 110 V (constant voltage) at 4 °C with standard Tris–glycine transfer buffer (25 mM Tris, 192 mM glycine, pH 8.3, 20% (v/v) methanol). Non-fat milk protein (1% in PBS) was used to block non-specific binding. Membranes were incubated with anti-NLRP3 or anti-β-actin for 16 hr at 4 °C. Membranes were washed extensively with PBS containing 1% (v) Tween-20, incubated for 1 h with HRP-conjugated secondary antibody, and developed with enhanced chemiluminescence (ECL) reagent. Protein immunoreactivity was visualized with an LAS4000 Gel Imager. Densitometric analysis was completed with ImageQuant software (GE Healthcare).

### 2.7 Detecting NLRP3 oxidation with NEM labelling

Reactive cysteine residues were labelled with NEM by reverse thiol trapping prior to analysis by mass spectrometry (LC-MS/MS). NLRP3-GFP was captured from the whole cell lysates of two 10-cm plates (∼3.0 mg total protein) using GFP-Trap resin (400 μL, 50:50 slurry) in PBS. The bead-bound NLRP3-GFP was incubated with chemical oxidant (10 μM CuCl_2_) or vehicle; the reaction was quenched with EDTA and sequentially treated with IAM and DTT as described above. After reduction, the bead samples were exposed to NEM (2 mM, 1 hr at 23 °C). Beads were washed twice with PBS and resuspended in SDS-PAGE sample buffer; the supernatants were collected using a mini-spin column and then proteins were separated by SDS-PAGE for subsequent excision and in-gel tryptic digestion of the NLRP3-GFP protein band. The resulting peptide solutions were stored at −20 °C for LC-MS/MS analysis.

### 2.8 In-gel tryptic digestion and LC-MS/MS analysis of isolated NLRP3-GFP protein

The ∼145 kDa Coomassie-stained band corresponding to NLRP3-GFP was excised from the SDS-PAGE gel and ‘in-gel’ digested with trypsin (2.5 μl of 0.5 μg/μl) at 37 °C for 16 h. The digest was acidified to 0.1% (v/v) formic acid with 4% (v/v) acetonitrile, and peptides were purified with C18 ZipTip (Millipore-Sigma). Tryptic peptides were analyzed with an Orbitrap Fusion Lumos Tribrid mass spectrometer that was coupled to an Easy-nLC 1200 liquid chromatography system and operated with Xcalibur (v4.0.21.10). Peptides were loaded in solvent A (0.1% (v/v) formic acid in H2O) onto a C18 trap column (75 μm x 2 cm; Acclaim PepMap 100, P/N 164946; ThermoScientific) at a flow rate of 2 μl/min. Trapped peptides were eluted onto a C18 analytical column (75 μm x 50 cm; PepMap RSLC C18; P/N ES803A; ThermoScientific) at a flow rate of 0.3 μL/min. The separation of peptides was completed using a linear gradient 5-28% of solvent B (0.1% (v/v) formic acid in 80% (v/v) acetonitrile) for 22 min followed by another linear increase to 40% solvent B over 3 min. Peptides were introduced into the ion transfer tube of the Orbitrap Lumos by electrospray (2.1 kV at 300 °C) in positive mode. A full MS scan at a resolution of 120000 FWHM was first performed to detect precursor ions having a m/z between 375 and 1575 and a +2 to +7 charge. The Orbitrap AGC (Auto Gain Control) and the maximum injection time were set to 4e5 and 50 ms, respectively. The Orbitrap was operated in top speed mode with a 3 sec cycle time for precursor selection. The most intense precursor ions presenting a peptidic isotopic profile and having an intensity threshold of at least 2e4 were isolated using the quadrupole (isolation window: 1.2 m/z) and fragmented using HCD (30% collision energy) in the ion routing multipole. The fragment ions (MS2) were analyzed at a resolution of 30000 with the AGC and the maximum injection time set to 5e4 and 60 ms, respectively. Dynamic exclusion was enabled for 30 sec to avoid acquiring precursor ions with a similar m/z (± 10 ppm). The raw MS data (*.raw) files were converted into Mascot Generic Format (*.mgf) using RawConverter (v1.1.0.18; The Scripps Research Institute) operating in a data dependent mode. The mgf file was used to search the SwissProt human protein database using Mascot algorithm (Matrix Sciences; version 2.7). Search parameters for MS data included trypsin as enzyme, a maximum number of missed cleavages of 1, charge state of +2 to +7, +57.021 for cysteine carbamidomethylation as fixed modification, +125.048 as a variable modification to identify NEM labelled peptides, methionine oxidation as variable modification, and a mass error tolerance of 10 ppm. A mass error tolerance of 0.02 Da was selected for the fragment ions. Only peptides identified with a score having a confidence higher than 95% were kept for further analysis. The resultant Mascot files were imported into Scaffold (v5.3.2, Proteome Software Inc) for comparison of different samples based on their mass spectral counting.

### 2.9 Identifying the reactive Cys landscape of NLRP3 in THP-1 cells

THP-1 monocytes were cultured in RPMI-1640 media supplemented with 2 mM glutamine and 10% FBS and then differentiated with PMA for 3 days, with media exchange after the first 24 h. Cells were resuspended in serum free media (∼18 h) prior to exposure to LPS (10 ng/mL, 4 h). After this, cells were washed into serum-free media lacking potassium (ΔK^+^). The pan-caspase inhibitor (ZVAD, 50 μM) was added 1 h prior to ΔK^+^ and maintained for the duration of the experiment After 1 h of ΔK^+^ media exposure, IAA-C6 (20 μM) was added for an additional 1 h incubation period. THP-1 cells were then collected and lyzed with RIPA buffer. The resulting whole cell lysates were clarified by brief centrifugation and then immediated used for the capture and identification of IAA-C6-labeled peptides by LC-MS/MS.

### 2.10 LC-MS/MS analysis of THP-1 cell lysates treated with IAA-C6

Proteins (1 mg) in whole cell lysates of THP-1 cells treated with IAA-C6 were reduced with DTT, alkylated with IAM and then adjusted to pH ∼8.0. Magnetic SP3 beads (5 mg) were added followed by immediate addition of 1 volume of 100% EtOH. Samples were vortexed and incubated for 10 min at RT with 1000 rpm agitation to maintain bead suspension. After protein binding, SP3 beads were washed extensively with 80% EtOH. Ethanol was removed and beads were allowed to air dry for 60 s prior to addition of Tris buffer (50 mM Tris, pH 8.0) and 20 μg of trypsin. Samples were digested overnight at 37 °C with 1000 rpm agitation. The peptide supernatant was collected and used for CuAAC ‘click chemistry’ reaction. Biotin-DADPS-Azide (light or heavy, 30 μM), BTTES (1 mM), CuSO_4_ (1 mM), and sodium ascorbate (15 mM) were added to each sample with vortexing after each addition. Solutions were incubated in the dark with end-over-end agitation for 90 min. Free biotin azide was removed by addition of alkyne-agarose (45 min). Samples were combined and biotin-linked peptides were captured on streptavidin-coated T1 Magbeads (400 μg per sample). After sequential washing with Tris buffer (containing 0.2 M Urea, 5% v/v ACN), PBS and water, peptides were eluted with two volumes of 5% v/v formic acid (30 min at RT with 1000 rpm agitation). The two elutions were combined and diluted with H_2_O to 2% v/v ACN, and dried in a SpeedVac before analysis by LC-MS/MS. Data were collected using a Vanquish Neo UHPLC coupled to an Orbitrap Astral mass spectrometer For each sample, peptides (∼1 µg) were captured on a PepMap Neo Trap column (C18, 300 μm x 5 mm, 5 μm particle size, 100 Å pore size, ThermoFisher Scientific), which was then submitted to reverse phase chromatography using an EASYSpray HPLC analytical column (75 μm x 50 cm, C18, 2 μm particle size, 100 Å pore size, ThermoFisher Scientific) that was subsequently developed at a flow rate of 250 nL/min at 40 °C using a multistep gradient (i.e., 2-30% B, 50 min; 30-45% B, 10 min) followed by a column wash (30 min). Mobile phase A consisted of 0.1% (v/v) formic acid in water, and mobile phase B consisted of 0.1% (v/v) formic acid in 80:20 acetonitrile:water (v/v). Spectra were acquired in positive mode using data-dependent acquisition (DDA) with a 1 second cycle time. Full MS scans (375-1875 m/z) were performed at 120,000 resolution (at m/z 200) with maximum injection time set at 50 ms and normalized AGC target at 200%. Precursors with charge states +2-8 were selectively isolated (1.6 Th quadrupole isolation window) for fragmentation using stepped HCD at Normalized Collision Energies 30%. Global source parameters included spray voltage at 1800V, ion transfer tube set to 275 °C and RF Lens at 35%; the Dynamic Exclusion window was set to 10 s. MS2 scans were performed using the Astral mass analyzer set at 150-2000 m/z, a maximum injection time of 10 ms, and AGC target of 300%. FragPipe was used to analyse spectra collected from whole cell lysates derived from THP-1 cells labeled with IAA-C6. The search routine was performed against the full proteome databases with the following parameters: cysteine carbamidomethylation (Δm = +57.021465 Da) as fixed modification, methionine oxidation (Δm = +15.9949Da) and N-termini acetylation (Δm = +42.0106Da) as variable modifications; 10 ppm for precursor mass tolerance; 20 ppm for fragment mass tolerance; trypsin as enzyme for protein digestion with 3 allowed missed cleavages, FDR of 1%. Quantification was performed based on MS1 intensity of isotopically-labelled peptides with Cys Δm of +280.1899 (light) and +282.2022 (heavy) as variable modifications in search parameters. A differential expression analysis was performed on the identified peptides using RStudio. Labelled (heavy/light) peptides were filtered for containing signal intensity in at least two replicates in one condition and the intensity values were log_2_ transformed. For missing values, an adjusted peruses style imputation method was applied with a 3.0 x standard deviation downshift and 0.2 width (variation factor). Peptide differential expression significance threshold was set at log_2_FC >2 and p < 0.05 (Benjamini Hochberg FDR adjusted).

### 2.11 Molecular dynamic (MD) simulations

Three adjacent NLRP3 units were extracted from the decameric NLRP3 cage structur*e* (PDB ID: 7PZC, [8]). The NACHT domain structures were removed, and a final trimeric LRR complex was subjected to MD simulations. No interchain NACHT-NACHT domain contacts were noted for 7PZC, so removal of this region was not expected to impact quaternary structure. CHARMM GUI was used to generate non-standard amino acid substitutions (CSO, Cys-SOH), of Chain A of 7PZC and to solvate the protein. To simulate the LRR interactions of the quaternary structure, high strength force fields were applied to the regions that would be interacting with neighboring LRR units. Additional positional restraints were applied to the N-terminal residues that were newly exposed due to deletion of the NACHT domain. GROMACS was installed locally and used to perform the MD production runs [30]. System energies were minimized with the steepest descent method and equilibrated using constant-substance, constant-volume, constant temperature ensemble (NVT) then constant-temperature, constant-pressure ensemble (NPT). The molecular systems were simulated for 60 ns, and downstream RMSD, RMSF, SASA (Solvent-Accessible Surface Area) and hydrogen bond occupancy were calculated with in-built GROMACS functions.

### 2.12 Data analyses

Data are presented as the mean ± S.E.M., with n indicating the number of independent, biological replicates. Statistical analyses and plots were completed with GraphPad Prism v6.0 or R Studio. The NLRP3 protein structures were visualized with ChimeraX (v1.9). Gemini AI was used to assist in code development for MD simulations with GROMACS and CHARMM-GUI.

## 3. RESULTS

### 3.1 NLRP3 is susceptible to Cys oxidation *in vitro*

The NLRP family as a whole, as well as NLRP3 specifically, display high Cys content relative to the human proteome [17]. There appears to be equitable distribution of Cys residues throughout the entire NLRP3 structure with many Cys side-chains observed to be solvent accessible. The reactive cysteine landscape of NLRP3 was initially assessed using a reverse thiol trapping approach [31, 32]. As shown in **Figure 1A**, NLRP3(WT)-GFP was captured using a GFP-Trap (i.e., an immobilized anti-GFP camelid nanobody) pull-down method. Bead-bound NLRP3(WT)-GFP was incubated in the absence or presence of Cu (II) chloride (1 μM CuCl_2_, 10 min), a well-described chemical oxidant that targets the thiol groups of Cys residues in proteins [33, 34]. After protecting native disulfides (S-S) and reduced Cys (-SH, thiol) residues with IAM, any oxidized Cys residues were reduced and marked by treatment with AMS or Mal-PEG5K. NLRP3(WT)-GFP was treated with increasing concentrations of CuCl_2_ (0.01 μM to 25 μM; **Figure 1B**), followed by sequential exposure to IAM, DTT and AMS. Low concentrations of CuCl_2_ (0.01-0.2 μM) with subsequent AMS labeling did not impair NLRP3 mobility during SDS-PAGE. A band shift was observed for CuCl_2_, ≥0.5-1 μM; the band shift was further accentuated under increased oxidation conditions (e.g., 10 μM CuCl_2_). Notable shifts in NLRP3 migration, representing the pool of reactive disulfides, were apparent between untreated and treated (IAM, DTT, and AMS). When NLRP3 was subjected to oxidant (10 μM CuCl_2)_ and treated with IAM and AMS only, then marginal retardation of NLRP3 migration was observed, indicating the dependency of DTT to restore oxidized Cys and reduce native disulfides. If IAM was omitted, then NLRP3 migration was slightly impaired. This suggests the occurrence of less reactive Cys residues that were not oxidized and were still available for reaction with IAM. Thus, it appears that the Cys oxidization of NLRP3 was titratable.

**Figure 1.**
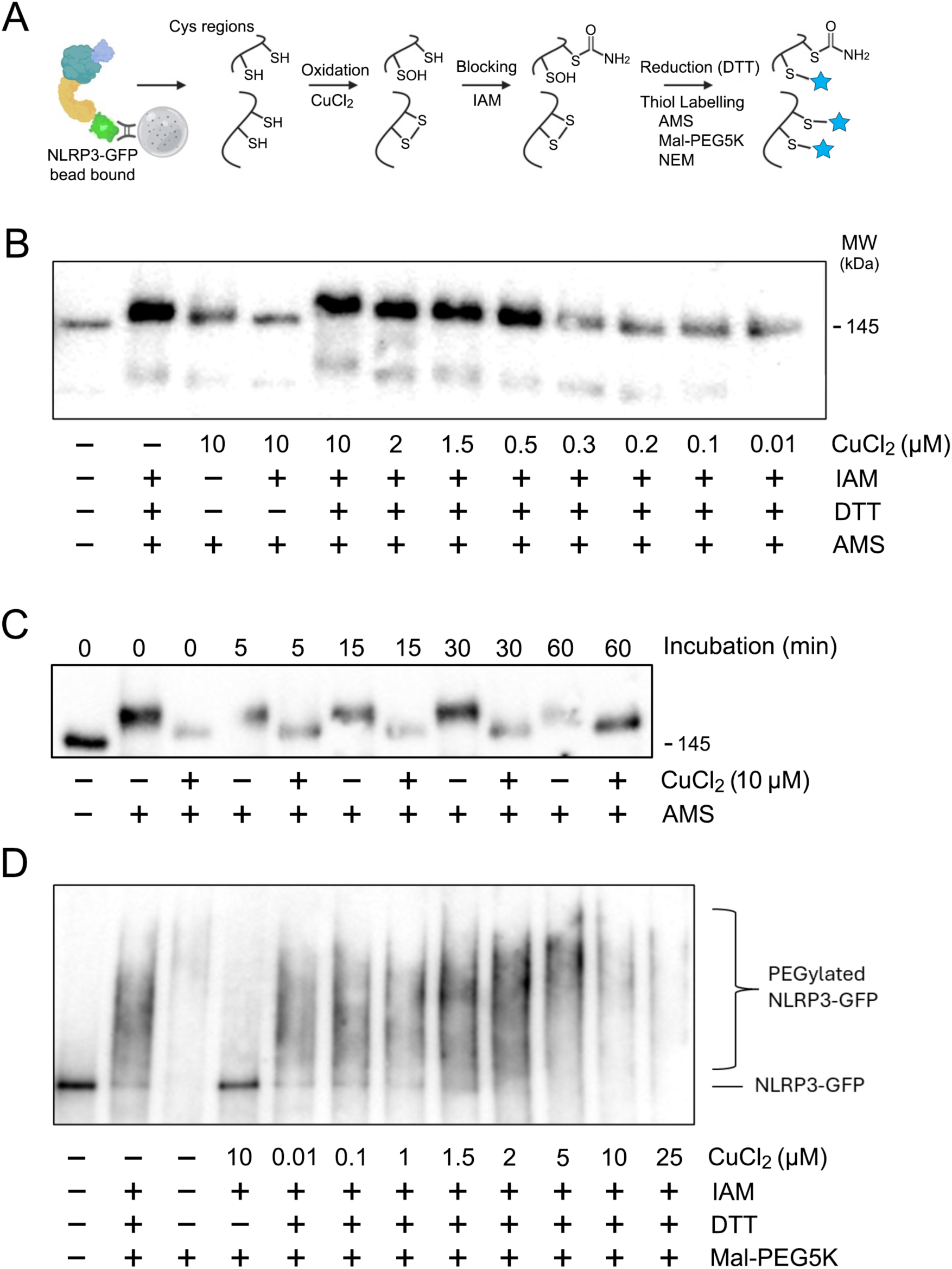
Profiling the reactive cysteine landscape of NLRP3. (**A**), Reaction scheme for labelling oxidizable Cys residues. NLRP3-GFP was overexpressed in HEK293T cells and then captured with GFP-Trap Sepharose resin. Bead-bound NLRP3-GFP was exposed to oxidant (CuCl_2_), remaining thiols were alkylated with iodoacetamide (IAM, 20 mM, 30 min, dark), subsequent DTT exposure (30 mM, 15 min) generated thiols of any oxidized Cys (including native disulfides) which were then reacted against a maleimide-based probe. (**B**), NLRP3-GFP was exposed to increasing chemical oxidant concentration (0 – 25 μM CuCl_2_); oxidized Cys residues were tagged with 4-acetamido-4’-maleimidylstilbene-2,2’-disulfonic acid (AMS) by reverse thiol trapping. (**C**), The stability of Cys oxidation events was assessed; NLRP3-GFP was captured on GFP-Trap beads and exposed to chemical oxidant (10 μM CuCl_2_; 10 min). After copper chelation with addition of EDTA, bead-bound NLRP3-GFP was buffer-exchanged (3 washes). Samples were held at room temperature for the indicated amount of time and then reacted with AMS. (**D**), NLRP3-GFP was exposed to 0 – 25 μM CuCl_2_; oxidized Cys residues were tagged with Maleimide-PEG_5000_ (Mal-PEG5K) by thiol trapping. Mobility shifts for NLRP3-GFP in (**B**) and (**D**) were assessed by SDS-PAGE and subsequent immunoblotting. Results are representative of two independent replicates.

The stability of the Cys oxidation was examined in a separate series of experiments. In this case, the chemical oxidant was removed by buffer exchange and then the NLRP3 protein samples were held in storage buffer for up to 60 minutes before direct thiol labelling with AMS. The reaction of NLRP3(WT)-GFP with AMS under control conditions (i.e., no CuCl_2)_ resulted in a notable retardation of gel migration (**Figure 1C**). This change in migration was generally negated with prior exposure of NLRP3 to 10 μM CuCl_2._ Some samples were treated with chemical oxidant, buffer-exchanged and then allowed to rest at room temperature prior to AMS reaction. Similar migration patterns were observed for NLRP3 when left for 5, 15 and 30 min before thiol trapping. These results imply that Cys residue modifications were still present, that is, the oxidized Cys residues were unable to react with AMS which in turn restored gel migration to levels observed in the control condition. With 60 min of waiting time, some loss in the oxidation-dependent inhibition of thiol labelling was observed. Moreover, the AMS reaction following 60 min incubation also appeared to be less effective under control conditions. This suggests some NLRP3 thiols became unavailable for reaction during an extended incubation period. Ultimately, the Cys oxidation events were judged to be stable for at least 30 min, and the Cys oxidation status of NLRP3(WT)-GFP did not revert spontaneously during the time required to complete the thiol trapping.

A similar sensitivity to CuCl_2 w_as observed when the final labelling of reactive Cys residues was completed with Mal-PEG5K (**Figure 1D**). Reaction with Mal-PEG5K in the absence of oxidative insult resulted in marked ‘smearing’ of the immunoreactive signal, with higher molecular weight NLRP3 species and decreased intensity of the unmodified NLRP3 band. The observation implied extensive PEGylation of NLRP3, with many Cys residues in NLRP3 maintained as thiols under basal conditions. Additionally, the native disulfide bond content was revealed under conditions that lacked chemical oxidant but did expose NLRP3 to IAM, DTT and Mal-PEG5K. In the absence of chemical oxidant, IAM treatment would alkylate all Cys residues except disulfide bonds. Subsequent exposure to DTT would reduce disulfide bonds and make them available for reaction with Mal-PEG5K. No qualitative difference in appearance was observed in the migration of control (no oxidation) and treatment samples (0.01-1 μM CuCl_2)_, which indicates no change in the reacted Cys content among these samples. The results imply that the same population of native disulfides in NLRP3 were accessible for Mal-PEG5K labelling for CuCl_2 u_p to 1 μM. It was only with higher oxidation potential (≥1.5 μM CuCl_2)_ that pronounced gel migration changes were observed for NLRP3. More extensive smearing with greater accumulation of high MW NLRP3 species and a decrease in the native NLRP3 band intensity were observed in samples exposed to 1.5-25 μM CuCl_2._ These results suggest novel Cys residues could be captured by the assay under conditions of elevated oxidation potential. It was also apparent that the total signal intensity was conspicuously reduced for the 10-25 μM CuCl_2 t_reatments. This was likely due to extensive PEGylation of NLRP3 which in turn prevented these species from entering the separating gel during SDS-PAGE.

When NLRP3 was oxidized but not treated with DTT, NLRP3 migration most closely resembled the untreated, control sample. This condition is best compared against the 10 μM CuCl_2 s_ample since they represent identical oxidative exposures (other than the inclusion of DTT). The absence of any significant PEGylation (absence of smearing with retention of native band intensity) indicates that the reaction conditions were sufficient with upstream oxidation and IAM-dependent thiol blocking affecting nearly all Cys residues. No thiols were generated in the absence of DTT prior to Mal-PEG5K labelling. Collectively, the results of AMS and Mal-PEG5K treatments reveal congruent phenomena. These results establish a titratable range of low (1 μM CuCl_2)_ and high (10 μM CuCl_2)_ oxidation events where some, but not all, Cys residues were oxidized.

### 3.2 Proteomic assessment of NLRP3 for the detection of reactive Cys residues

An experimental modification of the reverse thiol trapping method was coupled with mass spectrometry to identify the specific Cys residues of NLRP3 that were responsive to chemical oxidation. In this case, N-ethylmaleimide (NEM) was used as the terminal thiol label since it is easily identified with a distinctive mass shift during MS/MS fragmentation. Given the different pools of Cys reactivity observed with Mal-PEG5K and AMS labelling experiments, low (1 μM CuCl_2)_ and high (10 μM CuCl_2)_ chemical oxidant levels were used. On-bead digestions of captured NLRP3(WT)-GFP were initially completed, but these samples provided low-quality MS results. So, samples were subjected to SDS-PAGE, and a band corresponding to the MW of NLRP3(WT)-GFP was excised, trypsin-digested, and then analysed by LC-MS/MS.

Nine total replicates of the NEM labelling were subjected to LC-MS/MS analysis (two with high oxidant, five with low oxidant, and two under control conditions); 17 of the 45 Cys residues in NLRP3 were identified to contain the characteristic +125.05 Da mass that represents the addition of NEM (**Table 1**). The proteomic survey defined the reactive Cys landscape of NLRP3(WT)-GFP to include C38, C108, C150, C196, C279, C280, C514, C669, C673, C762, C770, C819, C843, C876, C884, C933, and C941. Many reactive Cys residues (12/17) were coincidently detected in the control sample (i.e., not treated with chemical oxidant). These NEM-labelled peptides may represent inherent disulfides; however, available structures of NLRP3 do not support the existence of disulfide bonds at these locations. There are a few possible explanations for the number of NEM-labelled cysteines detected in the control samples. The reverse thiol trapping protocol modified nearly all reactive Cys prior to DTT reduction and NEM-labelling based on western blot observations (i.e., AMS or Mal-PEG5K; **Figure 1B,D**). Detection of modified Cys residues by mass spectrometry is sensitive, and it could be expected that some Cys residues escaped blocking and were labelled with NEM. This likely accounts for some of the detected Cys in the negative control. Alternatively, treatment with DTT after blocking could destabilize the structure of NLRP3 and expose some buried residues, enabling reaction with NEM in subsequent steps. And finally, Cys residues could be oxidized during cell lysis and sample processing. Overall, it is moderately surprising how many of the reactive Cys residues were detected as NEM modified in the absence of oxidation by MS, when western blotting revealed a limited pool of maleimide-reactive sites in the control samples. Nonetheless, the Cys residues modified by NEM across most replicates and conditions represent reactive cysteines that likely support NLRP3 function.

**Table 1.**
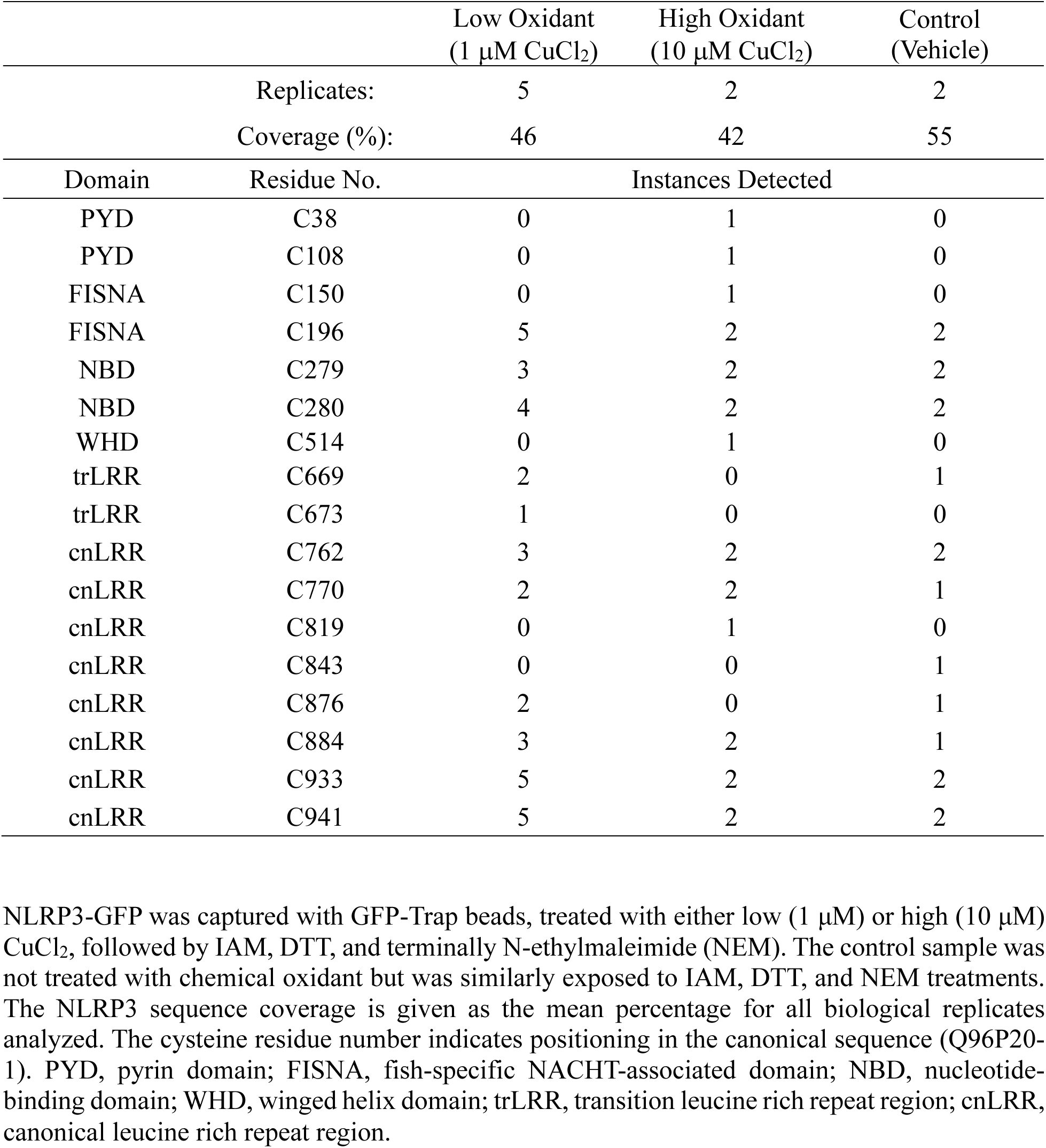
Reactive Cysteine Landscape of NLRP3. NLRP3-GFP was captured with GFP-Trap beads, treated with either low (1 μM) or high (10 μM) CuCl_2_, followed by IAM, DTT, and terminally N-ethylmaleimide (NEM). The control sample was not treated with chemical oxidant but was similarly exposed to IAM, DTT, and NEM treatments. The NLRP3 sequence coverage is given as the mean percentage for all biological replicates analyzed. The cysteine residue number indicates positioning in the canonical sequence (Q96P20-1). PYD, pyrin domain; FISNA, fish-specific NACHT-associated domain; NBD, nucleotide-binding domain; WHD, winged helix domain; trLRR, transition leucine rich repeat region; cnLRR, canonical leucine rich repeat region.

Notably, six Cys residues were determined to be uniquely modified with NEM following NLRP3(WT)-GFP exposure to chemical oxidant. These residues include C38, C108, C150, C514, C673, and C819. Of these residues, only C673 was uniquely detected with low oxidation treatment, potentially suggesting its high reactivity. Moreover, another 21 Cys residues were determined to be IAM modified across all replicates. Seven of these were uniquely modified by IAM and never detected with subsequent NEM reaction (including C261, C633, C837, C838, C843, C990, and C998). These residues represent Cys that exist as free thiols (-SH), are not native disulfides, and are not susceptible to oxidation. The residues may be internally located or otherwise unreactive due to constraints of the local microenvironment.

In a subsequent set of experiments, THP-1 macrophages were treated with LPS and then subjected to extracellular K^+^ deficiency (ΔK^+^) to activate the NLRP3 inflammasome. The cells were subsequently exposed to cell-permeable iodoacetamide derivative (i.e., stable-isotope labeled IAA-C6) after treatments with LPS (light IAA-C6) or LPS and ΔK^+^ media (heavy IAA-C6) to profile reactive cysteines *in situ* during inflammasome activation. Following proteomic processing, a total of seven IAA-C6-modified Cys peptides from NLRP3 were identified (**Table 2**). Two of the detected Cys residues are located in the FISNA (i.e., C150 and C196), and the remaining five are dispersed throughout the LRR (i.e., C770, C819, C876, C884 and C941). All reactive Cys residues were previously identified by reverse thiol-trapping following GFP-Trap purification of recombinant NLRP3-GFP protein and exposure to chemical oxidant (**Table 1**). All but one of the Cys residues were marked under the LPS stimulatory condition, that is the canonical “Signal 1” that is typically used as the transcriptional priming event for *NLRP3* and *IL1β* genes [3, 17]. The C941 residue was unique in that it was only marked after ΔK^+^ media treatment, whereas C819 and C876 were not detected after exposure to ΔK^+^ media.

**Table 2.** Reactive Cys profiling of NLRP3 in THP-1 macrophages with K^+^-depletion. IAA-C6 labeled peptides were recovered from THP-1-derived macrophages and characterized by LC-MS/MS. Cells were primed with LPS (4 h) and treated with potassium-deficient media (ΔK^+^) for 1h, followed by 1 h treatment with ΔK^+^ with 20 μM IAA-C6. The modified Cys (C*) residue number indicates positioning in the canonical sequence (Q96P20-1). Log2FC, log_2_ fold-change for LPS vs. ΔK^+^ treatments; FISNA, fish-specific NACHT-associated domain; LRR, leucine rich repeat region. Data represent 3 biological replicates for LPS and ΔK^+^; the incidence of peptide detection (/3) is provided. The incidence of peptide detection in the reverse thiol-trapping of recombinant NLRP3-GFP protein is provided in the last column with (#) of independent observances.

| Domain | Peptide Sequence | Modified Cys | LPS (/3) | ΔK<sup>+</sup> (/3) | Log2FC | ID’d in Thiol Trapping |
|---|---|---|---|---|---|---|
| FISNA | FQ**C\***IEDR | 150 | 3 | 3 | -0.46 | Yes (1) |
|  | T**C\***ESPVSPIK | 196 | 3 | 2 | -1.79 | Yes (9) |
| LRR | VLCETLQHPG**C\***NIR | 770 | 1 | 1 | - | Yes (5) |
|  | LL**C\***VGLK | 819 | 1 | 0 | - | Yes (1) |
|  | LYVGENALGDSGVAIL**C\***EK | 876 | 1 | 0 | - | Yes (3) |
|  | NPQ**C\***NLQK | 884 | 3 | 2 | -1.27 | Yes (6) |
|  | LLCEGLLHPD**C\***K | 941 | 0 | 1 | - | Yes (9) |

### 3.3 Positioning of reactive Cys residues in functional domains of NLRP3

As illustrated in **Figure 2A**, reactive Cys residues (i.e., possessing NEM modification subsequent to Cu^2+^ exposure) were localized to all functional domains of NLRP3 as well as some loops and linker regions that connect the various domains. Two reactive Cys residues, C38 and C108, within the PYD were identified under high oxidant conditions. C38 exists in a solvent exposed loop connecting a-helix 2 and 3 of the PYD (**Figure 2B**); a region that is important for NLRP3^PYD^ filament and ASC speck formation [35, 36]. C108 forms a disulfide bond with C8 [35], and a previous report found the C8 and C108 residues to be involved in agonist sensing during inflammasome activation [9]. Within the NACHT domain, reactive Cys residues were identified within the FISNA (C150, C196), NBD (C279, C280), and WHD (C514) regions (**Figure 2C**). Finally, the majority (10/17) of reactive Cys residues were identified throughout the LRR domain of NLRP3 (**Figure 2D**). The LRR can be partitioned into transition (trLRR, an important region for NEK7 recruitment [37]) and canonical (cnLRR, provides stabilizing contacts for homo-oligomerization regions [8, 14, 38]). Two reactive Cys residues were located in the trLRR (C669 and C673), and several were resolved to the cnLRR (C762, C770, C819, C843, C876, C884, C933, and C941).

**Figure 2.**
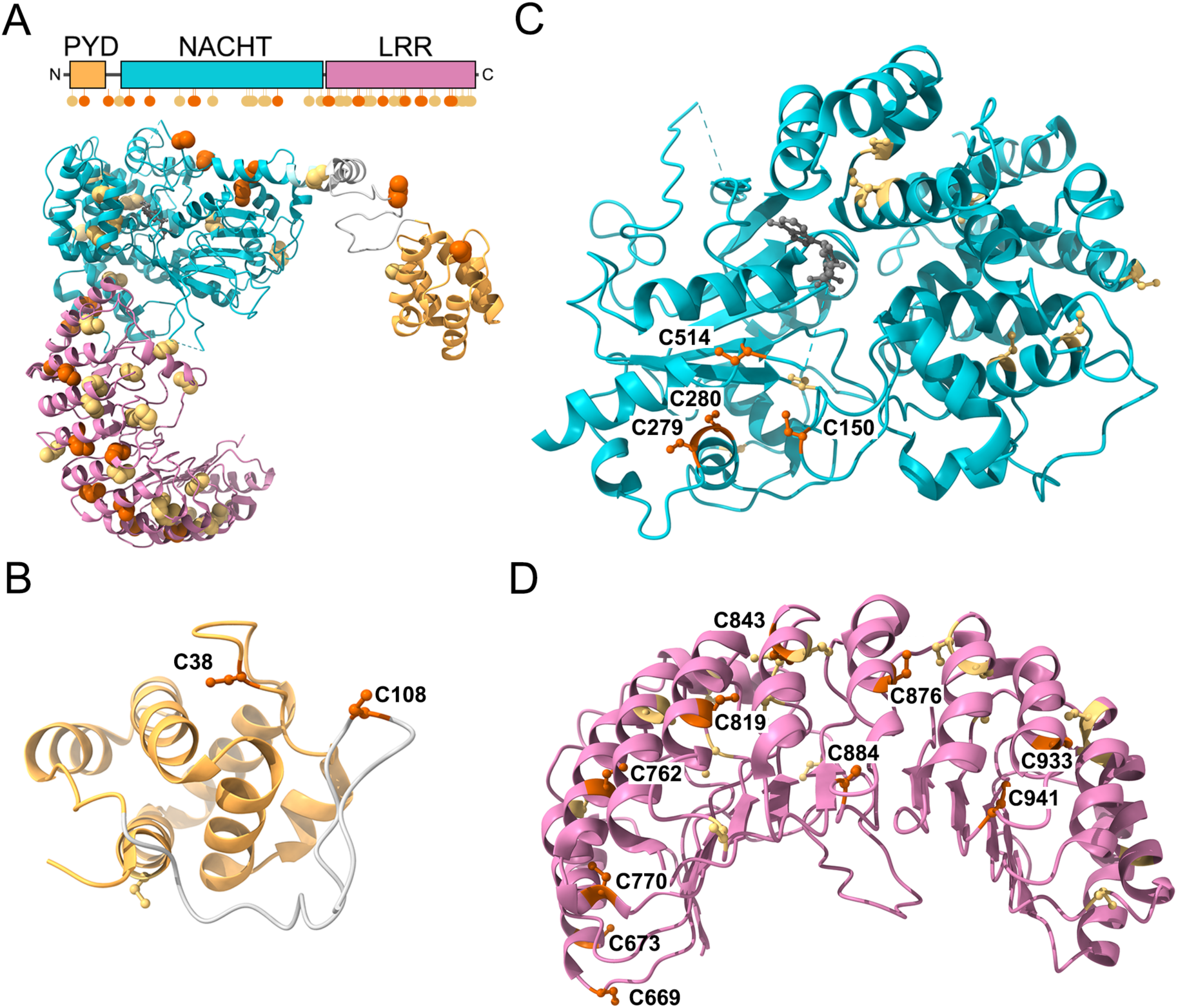
Positioning of reactive Cys residues on the NLRP3 protein. (**A**), Reactive Cys residues were mapped on the structure of full-length human NLRP3 (derived from the inactive human decamer ring; PDB ID: 7PZC). Red spheres represent reactive Cys sidechains that contained NEM labels; yellow spheres represent Cys sidechains that were not found to have NEM modification (only with IAM or not detected at all). The reactive Cys landscape of important NLRP3 protein domains is also provided: (**B**) pyrin domain, PYD (in orange); (**C**) <u>NA</u>IP, <u>C</u>IITA, <u>H</u>ET-E and <u>T</u>P1 domain, NACHT (in teal); and (**D**) the leucine rich repeat domain, LRR (in pink). The bound ADP molecule (gray) is shown in the nucleotide-binding domain. All structures were generated using ChimeraX.

### 3.4 Cys oxidation destabilizes LRR-LRR interactions in the inactive NLRP3

As illustrated in **Figure 3A**, the inactive NLRP3 oligomer (7PZC) assumes a cage-like structure that sequesters the N-terminal PYDs within the central core. Critical to the maintenance of the inactive cage are three distinct LRR-LRR interfaces. Of note, an acidic loop in the trLRR region provides the intermolecular contacts in the concavities of the two interacting LRR domains, and a separate LRR-LRR interface bridges the convex surfaces of adjacent NLRP3 proteins [8]. MD simulations were used to test the hypothesis that Cys oxidation of the LRR can elicit conformational rearrangements to destabilize the inactive quaternary structure. To generate the oxidized model, multiple Cys residues (i.e., C669, C673, C762, C770, C819, C843, C876, C884, C933, and C941) were modified into sulfenic acid (Cys-SOH) only in the central chain A of a trimeric LRR complex. Over the course of a 60 ns simulation, the native state structure reached equilibrium within 30 ns, while the oxidized model still displayed gradual increase in RMSD in the final 30 ns of simulation with higher overall RMSD observed throughout the production run (**Figure 3B**). The final native and oxidized structures were aligned on the central LRR of the trimer (chain A); harmonious overlap was observed for the C-terminal LRR region with divergent alignment apparent in the N-terminal segment. RMSD was elevated (4.92 Å) when all atom pairs were considered (**Figure 3C**). Some alignment disparities were also present in adjacent chains B and C of the trimer, when aligned on chain A. Taken together, these observations suggest the oxidized model maintained a less stable conformation.

**Figure 3.**
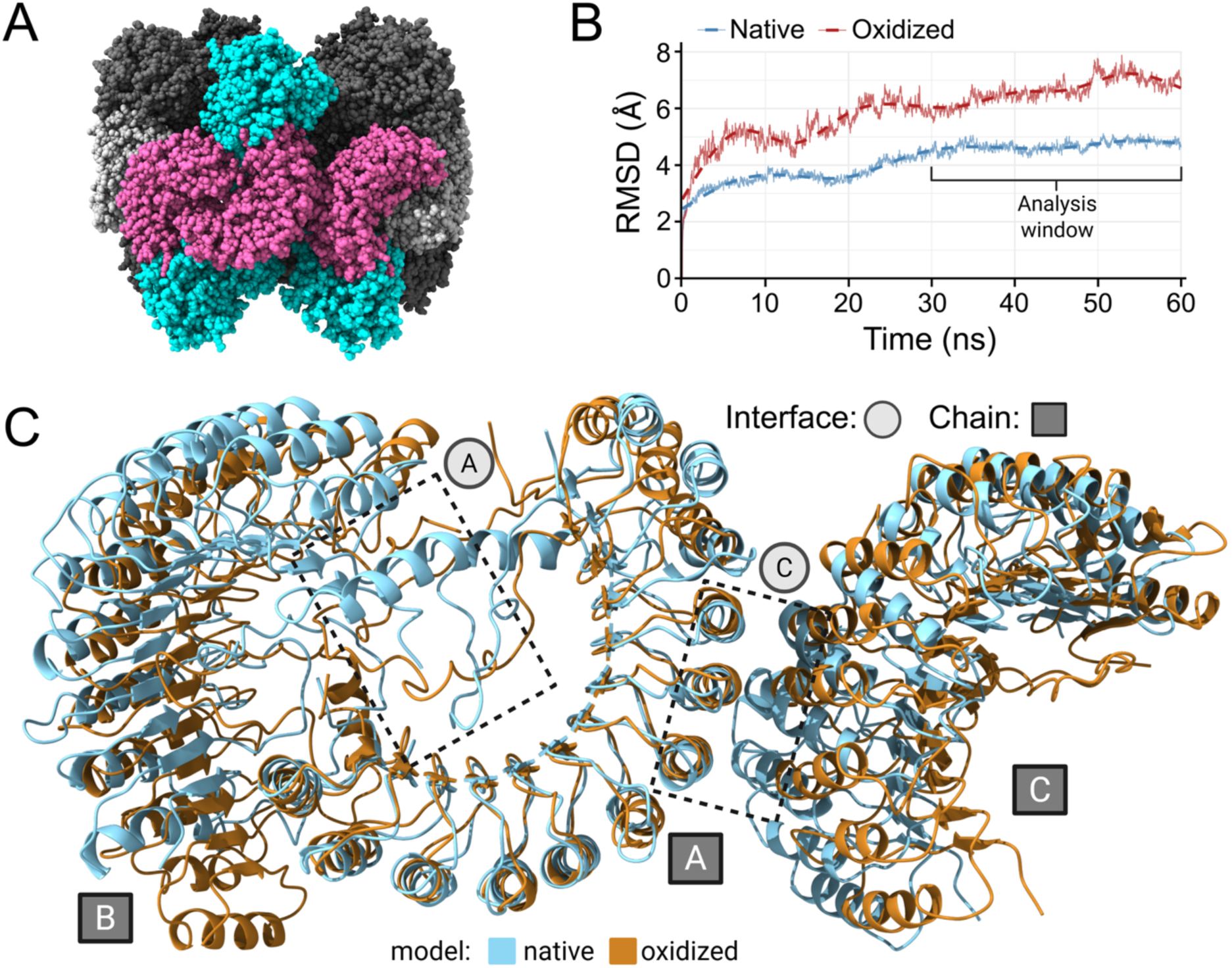
MD simulation models show the impact of oxidized Cys-SOH on the inactive LRR conformation. (**A**), A trimer of LRR domains (LRR, aa 649-1036, shown in pink. Chains A-C of PDB ID: 7PZC.) was extracted from the inactive, oligomeric NLRP3 structure. C669, C673, C762, C770, C819, C843, C876, C884, C933, and C941 were modified to sulfenic acid (Cys-SOH) in Chain A of the oxidized model. (**B**), Root-mean square deviation (RMSD) was calculated over the production run following a least-squares (LSQ) fit to the equilibrated reference structures at time 0 and plotted as a function of time (0-60 ns). (**C**), The final native (blue) and oxidized Cys-SOH (brown) models were aligned on the central Chain A. Interchain LRR-LRR interaction regions (interfaces A and C, as described in [8]) are outlined with dashed boxes. All structures were generated using ChimeraX.

The positioning and orientation of the trLRR fold, including the acidic loop of interface A that stabilizes the inactive cage, were more closely examined. The final topologies were generally similar for the native simulation model and the original cryo-EM structure following the 60 ns production run (**Supplemental Figure S1**). However, the upstream α-helix (α_L, a_a 707-719) was repositioned, reflecting an inherent flexibility in this region. In the Cys oxidation model, the trLRR region sampled an extensive conformational ensemble. Marked structural deviation was observed for α_L a_nd the acidic loop during the production run (**Figure 4A, panels i-iv**). At the beginning of the simulation, the acidic loop had similar positioning and interchain contacts as the native structure. At 3 ns, α_L w_as largely unfolded; by 19 ns, the helix was completely disordered and began to establish stable interactions with the LLR C-terminus of chain B, sites that were initially occupied by the acidic loop. As a result, the acidic loop was also significantly repositioned as the simulation progressed. Novel stabilizing contacts were established at 30 ns, including α_L r_esidues V704, S711, S712, H713, A714, A715 and C716 of chain A with the V1031’, E1033’ and W1036’ C-terminal residues of chain B. This novel topology was generally maintained for the remainder of the simulation, as seen in representative structures sampled at 30 and 60 ns (**Figures 4B** and **4C**, respectively). However, π-π stacking of H685 with W1036’ between Chain A and B residues was lost at 60 ns. S711, S712 and C716 contacts were present in both 30 and 60 ns frames, indicating their stability and presumed significant contribution to the novel interface. Hydrophobic interactions were also abundant with two valine (V704, V707) and two alanine (A714, A715) residues in the unfolded α_L a_nd a valine in the C-terminus of Chain B (V1031’). The molecular rearrangements to form a novel intramolecular interface occurred only in the trLRR for the oxidized chain A, and not in the adjacent B and C chains (**Supplemental Figure S2**). Moreover, the secondary helical structure of α_L w_as also maintained in non-oxidized chains B and C, and the radial geometry derived from the original structure was well preserved in the simulated LRR trimer model. The C chain α_L p_ositioning and acidic loop architecture were somewhat divergent; however, this was expected since chain C lacks a partnering LRR in the model to form interface A contacts. No significant structural alteration was observed during the simulation for interface C at the convex surfaces of A and C chains. Finally, the molecular rearrangements observed during the simulation were associated with a rapid loss in buried surface area for the acidic loop (**Figure 4D**). Coincidently, a marked increase in buried surface area (nearly +200 Å^2^) was predicted for the unfolded αL as it replaced the acidic loop in this molecular space. As the simulation progressed, more α_L r_esidues established contacts with a further increase in the buried surface area of this interface (to 300 Å^2^). Ultimately, the total buried surface area predicted for the trLRR loop in the Cys oxidation model recovered to occupy a space similar to the native structure (∼475 Å^2^).

**Figure 4.**
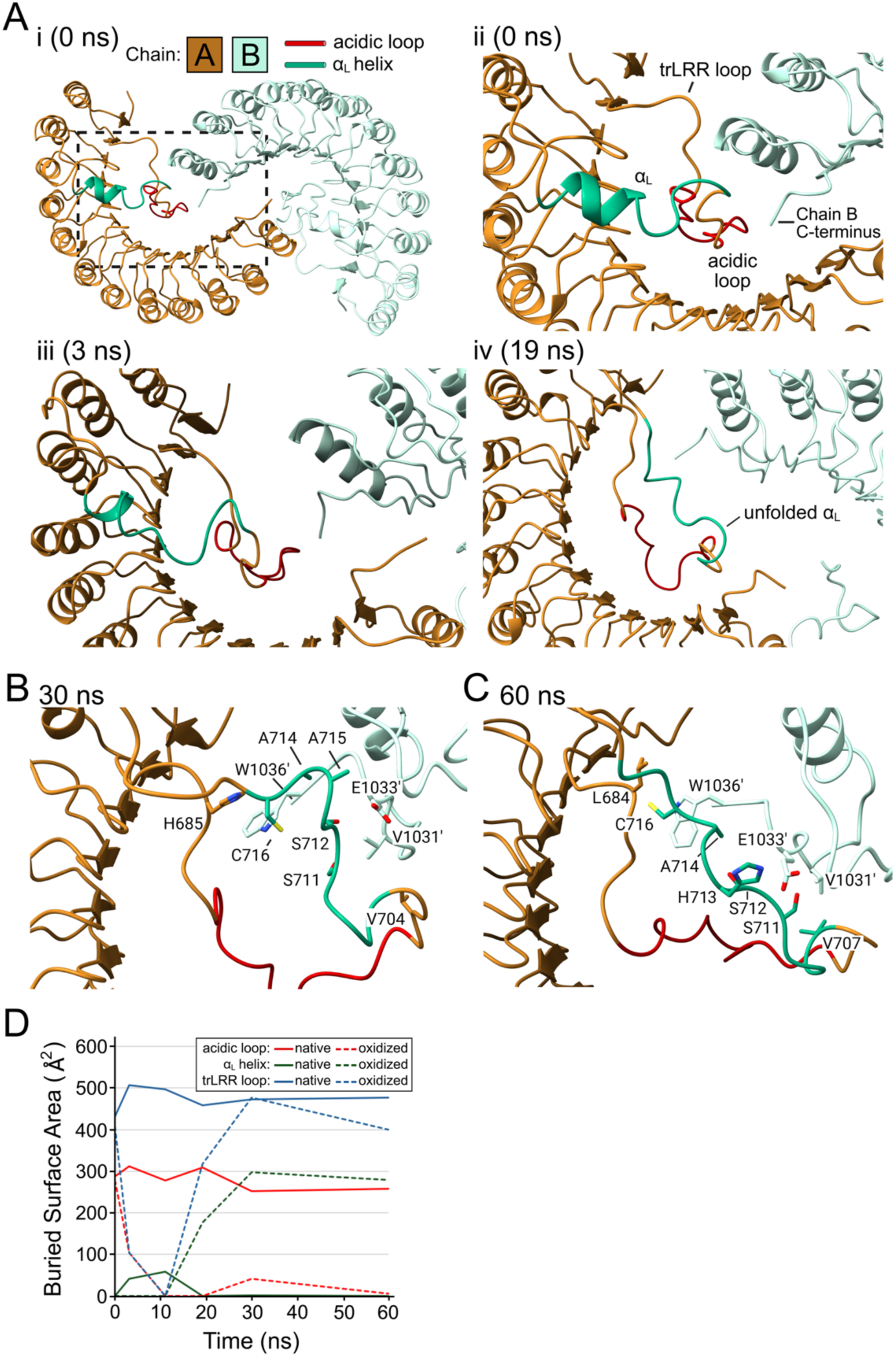
Structural dynamics of the trLRR loop in the Cys-SOH oxidation model during MD simulation. (**A**), The interchain LRR-LRR region (*panel i*, interface A described in [8]) is shown for the Cys-SOH oxidized model. As described in Materials and Methods, ten reactive Cys residues were modified to sulfenic acid in Chain A of 7PZC. Dynamic changes in the trLRR structure were tracked for the oxidized Cys-SOH model during the MD simulation. The starting structure (0 ns, *panel ii*) was captured prior to the first frame of the simulation but after the energy minimization step. At 3 ns (*panel iii*), the α_L_ helix was partially unfolded; at 19 ns (*panel iv*), the α_L_ helix was completely unfolded with extension of the acidic loop. Representative structures of the trLRR are shown at 30 and 60 ns (**B** and **C**, respectively) with novel contacts developed for the unfolded α_L_ helix (Chain A) and the LLR C-terminus (Chain B). All structures were generated using ChimeraX. (**D**), The buried surface area (Å^2^) was calculated for the trLRR region as well as the internal motifs (i.e., α_L_ helix and acidic loop) during the 60 ns MD simulation using ChimeraX.

Several analyses were performed to characterize the structural changes observed between the native and oxidized LRR models. There was increased RMSF in regions of the oxidized structure, including the acidic loop and α_L o_f interface A as well as interface C residues at the C-terminus (**Figure 5A**). Changes in the solvent accessible surface area (SASA) of several acidic loop residues (i.e., increased: K689, E690, E691, R699, and L701; decreased K696) were apparent (**Figure 5B**), but only modest changes in SASA changes were observed throughout interface C and the N-terminal interface A residues. Several interchain hydrogen bonds were lost from the oxidized structure, which agrees with a general destabilization of the acidic loop interface (**Supplemental Figure S3)**. As anticipated, modification of Cys residues to sulfenic acid yielded many changes in their local environments (**Figure 5C**). Oxidation of the reactive Cys residues disrupted the stable, very localized hydrogen bonding network in the native model and elicited other novel hydrogen bonds (**Figure 5D**; **Supplemental Figure S3**). Most of these novel bonds were transient, indicating the development of dynamic and unstable interactions (**Supplemental Figure S4**). C770, C884, C933 and C941 had the largest changes in SASA and intrachain hydrogen bonding which suggests these residues are the primary drivers of structural alterations. Notably, these residues were also reliably characterized as reactive during *in vitro* and *in situ* experimentation (**Tables 1** and **2**). The C770 residue, which gained the most SASA, also acquired several new hydrogen bonding partners, including R773, L737 and S740 (**Supplemental Figure S4**). Interestingly, bonds with L737 and S740 were mediated by the novel sulfene sidechain of C770. L737 is in the first LRR of the cnLRR, and S740 is C-terminal of the α_L (_aa 706-716). Thus, repositioning of S740 may be a significant driver in the overall restructuring of the trLRR upon oxidation, including the unfolding of the α_L h_elix and the destabilization of the acidic loop. Collectively, the MD simulation revealed the sulfenylation of 10 Cys residues across the LRR sufficient to destabilize the trLRR loop that is described to act as an autoinhibitory “lock” to secure interactions of concave surfaces in adjacent LRRs and to prevent premature inflammasome activation.

**Figure 5.**
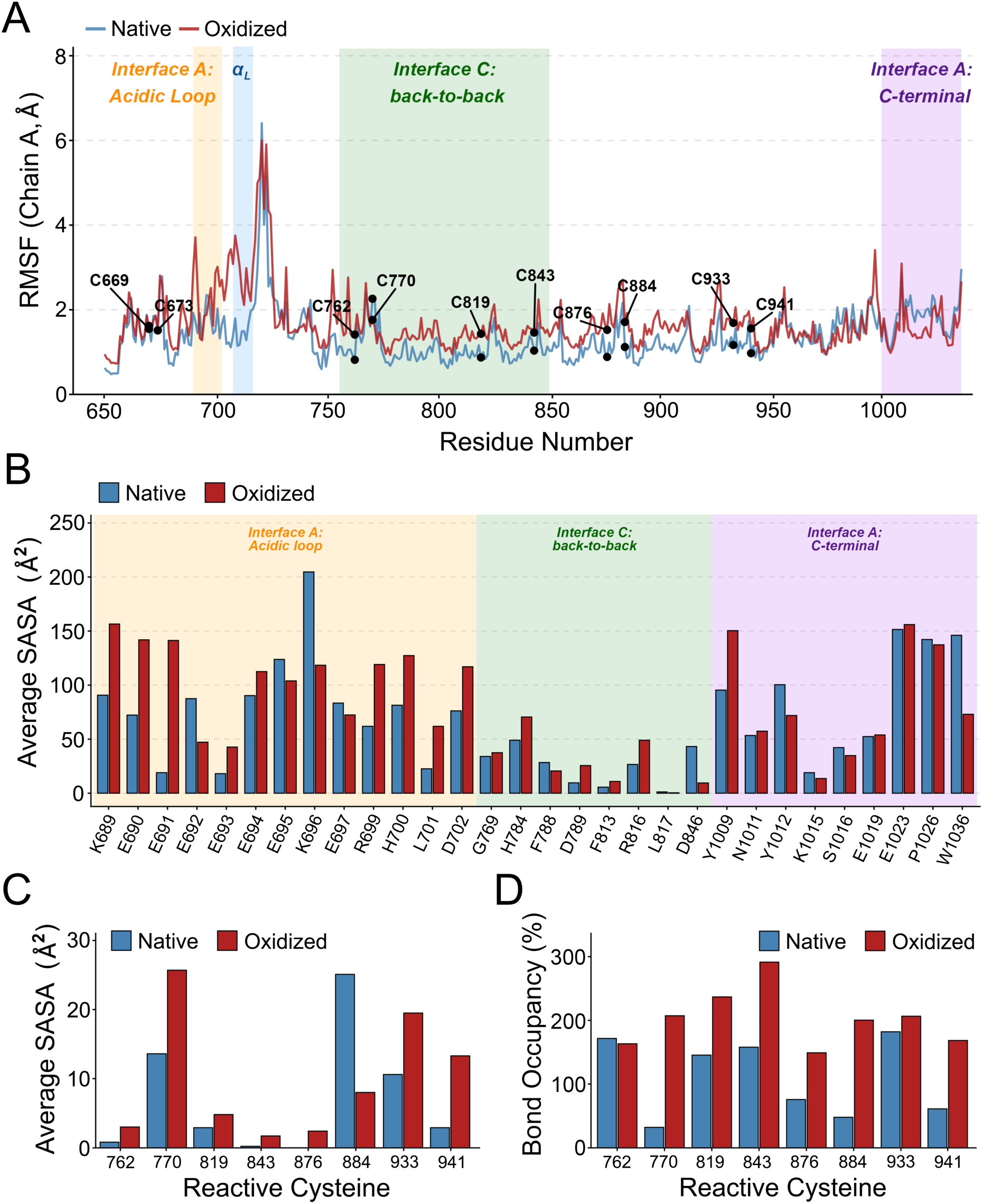
Impact of Cys-SOH oxidation on LRR-LRR interfaces and oxidized reactive cysteine residues. Root-mean square fluctuations (RMSF, **A**) and the solvent accessible surface areas (SASA, **B**) for each residue of chain A (LRR, aa 649-1036) are provided for the native (blue) and Cys-SOH oxidized (red) models. Average RMSF and SASA values were generated from the 30-60 ns window of the MD simulation. Important contact regions of the LRR are highlighted: interface A including the acidic loop (orange), α_L_ helix (blue), and C-terminal contacts (purple) as well as interface C (green). The average SASA (**C**) and cumulative hydrogen bonding occupancy (**D**) of reactive Cys residues were also generated for the native and oxidized models. SASA and hydrogen bonding occupancies were calculated in GROMACS.

### 3.5 Locational impact of reactive Cys residues in the NACHT domain

NLRP3 undergoes significant conformational changes upon activation with the ATPase activity of the central NACHT domain providing the biochemical switch to initiate the structural rerrangements required for oligomerization and assembly of the inflammasome complex [6, 10, 14, 39]. Four reactive Cys residues (i.e., C150, C279, C280 and C514) were identified (**Figure 2C**, **Tables 1 & 2)** and localized to important regions of the NACHT domain that are known to participate in molecular rearrangements of NLRP3 during the activation process [6, 39]. The reactive C150 and C196 residues reside in the FISNA subdomain; C279 and C280 are located in the NBD while C514 is a part of the WHD.

C150 is located in the FISNA subdomain at the N-terminus of loop 1 (**Figure 6A**). The FISNA provides a flexible loop to bridge the NACHT and PYD domains, and loop 1 transitions from a partially disordered state in the inactive NLRP3 structures [8, 37] to a fully ordered state that stabilizes the active NLRP3 disc [14]. Although the C150 and C514 sidechains are located in distinct subregions of the NACHT, both appear to coordinate with other residues in loop 1 of FISNA to stabilize the inactive structure (**Figure 6B**). The C150 positioning is coordinated in the inactive form by interactions with Q149 (sidechain) as well as K153 and R154 (backbone). In addition, the C514 residue, which is located in a hairpin β-loop of the WHD, coordinates with the Y143 sidechain in the loop 1 region of FISNA. The reactive C280, located in the NBD, coordinates with Y136 at the end of loop 1 in FISNA. The analysis of two available inactive NLRP3 structures (i.e., 6NPY and 7PZC) show the reactive Cys residues occupy similar positioning, regardless of NEK7 licencing (**Supplemental Figure S5**). The C196 sidechain was not resolved in either of the inactive NLRP3 structures solved by Sharif et al. (6NPY, [37]) or Hochheiser et al. (7PZC, [8]) due to disorder in this region of the NACHT domain.

**Figure 6.**
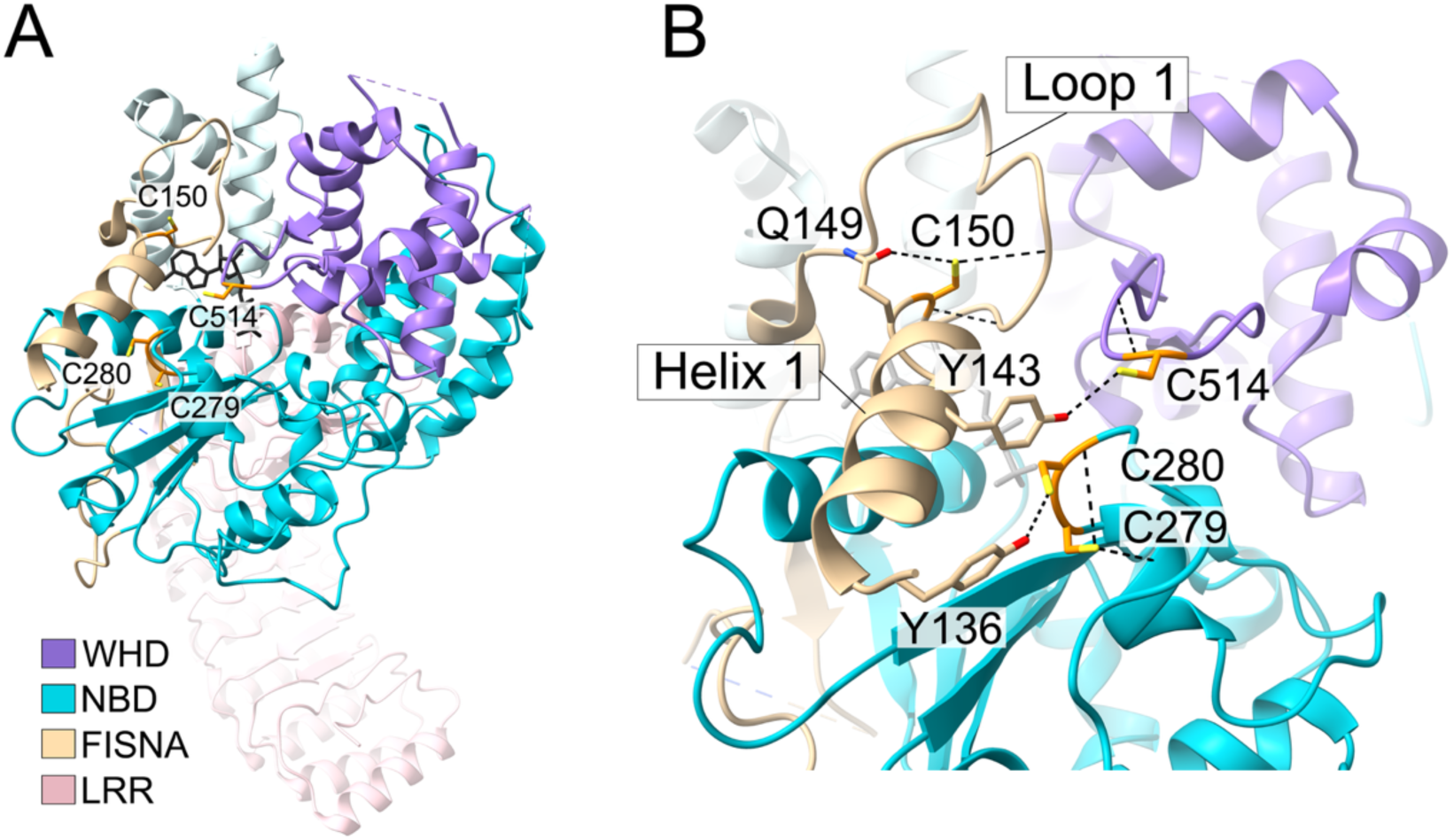
Reactive Cys residues are localized within the FISNA domain and are suggested to stabilize the inactive NLRP3 conformation. The inactive NLRP3 structure (ADP-ligated) was extracted from the NLRP3-NEK7 binary complex (PDB ID: 6NPY). In (**A**), the positioning of C150, C279, C280 and C514 residues are shown relative to the FISNA (beige), NBD (teal), WHD (purple) and LRR (pink) domains. The ADP molecule is shown in black, and the NEK7 structure was removed for clarity. In (**B**), molecular contacts of Cys residues (C150, C279, C280 and C514) are shown within core elements of the FISNA domain. All structures were generated using ChimeraX.

### 3.6 Functional impact of reactive Cys residues with NLRP3 inflammasome reconstitution

Individual mutations of reactive Cys residues were generated to evaluate their contribution to NLRP3 inflammasome activity (**Figure 7**). The impact of these mutations were assessed by reconstitution of the inflammasome in HEK293T cells (i.e., transient transfection of inflammasome components NLRP3, ASC, pro-caspase-1, and pro-IL-1β). After overnight transfection, IL-1β release was measured from the cell culture supernatant following exposure to nigericin. As expected, nigericin treatment was found to significantly increase IL-1β release from cells transfected with NLRP3(WT). Several NLRP3 single Cys mutants yielded reduced IL-1β secretion when compared against the WT inflammasome. The C150A (p < 0.05) and C196A (p < 0.01) mutations within the NACHT domain suppressed IL-1β by ∼ 25% whereas C514A had no effect. Two reactive Cys residues in the LRR, C933A (p < 0.05) and C941A (p < 0.05), were also found to significantly attenuate IL-1β secretion. In agreement with previous studies [9], the single C108A mutation did not have significant impact on IL-1β secretion.

**Figure 7.**
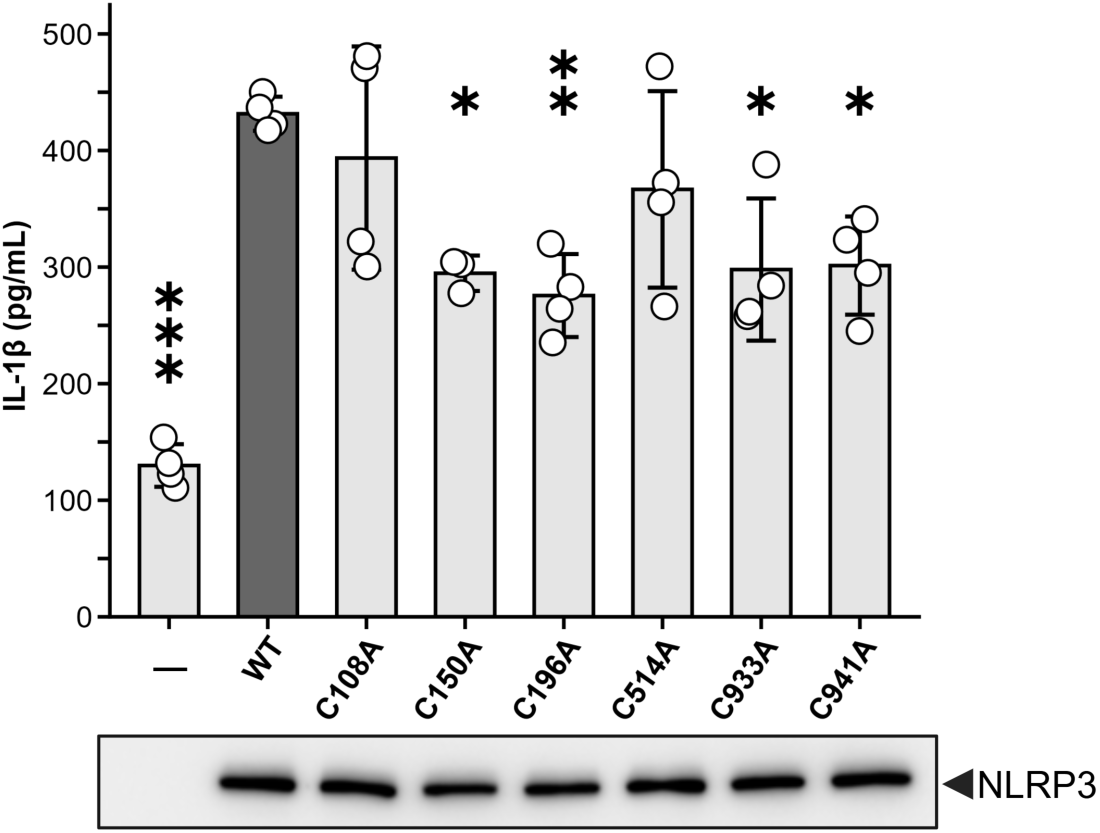
Impact of Cys mutation on the NLRP3 response to nigericin. The impact of Cys mutation) on NLRP3-dependent IL-1β maturation following inflammasome reconstitution in HEK293T cells. At 18 h post-transduction with ASC, pro-caspase-1, pro-IL-1β and NLRP3 (WT or C/A mutants), cells were treated with 10 μM nigericin (1 h). Secreted IL-1β was measured by ELISA and normalized to WT levels within each replicate. Western blots of FLAG-tagged NLRP3(WT) and the Cys mutants from whole cell lysates show comparable protein levels. Significantly different from WT, one-way block ANOVA of normalized means with Dunnett’s post-hoc test: *-p< 0.05; ** p < 0.01, ***-p < 0.001.

## 4. DISCUSSION

The dependency for ROS signaling during NLRP3 inflammasome activation is well established [16, 17]. However, the precise role played by these physiological signalling molecules in the licensing and activation of the NLRP3 inflammasome has remained largely unexplored. Modern redox biology has advanced that ROS are not simply toxic biproducts of dysregulated homeostasis, rather ROS are integral signalling molecules produced by cells to mediate diverse responses [20, 21]. In line with this paradigm, we expected NLRP3 to contain ROS-sensitive Cys residues that would confer functional impacts when subjected to post-translational (oxidative) modification. The present report provides the first exploration for such residues in NLRP3 and further establishes their functional significance in the process of inflammasome activation.

NLRP3 appears to possess hierarchical *in vitro* sensitivity to oxidative signals since Cys residues of NLRP3 were categorized by reverse thiol trapping into two general populations that reflect more and less thiol reactivity. The abundance of reactive Cys residues in no oxidation controls was unexpected since native disulfides would be labelled in the reverse thiol trapping approach. While it is possible that these Cys form disulfides as part of a transient structure that has not yet been determined, current structural data suggest it is unlikely, if not impossible that these Cys residues form disulfides within NLRP3 (intra or intermolecular). Therefore, the detection of these liganded Cys in the control samples suggests either incomplete chemistry during thiol trapping or oxidation during to sample processing. IAA-C6 was applied *in cellulo* to THP-1-derived macrophages undergoing inflammasome activation to validate the reactivity of the Cys residues. A total of 7 Cys sites were liganded, all of which were identified with *in vitro* experiments. The identification of reactive cysteines (e.g., C150 and C196 in the FISNA) from a native biological context validates the residues identified from *in vitro* experimentation, since the protein context and therefore cysteine reactivity in the cell varies drastically from a purified, bead bound state.

Most (8/10) of the reactive Cys in the LRR are present in 10 or more of the 14 NLRP family members when considering a previous sequence alignment analysis [17]. In contrast, the reactive Cys residues in other domains were minimally conserved in the family; the residues identified in PYD (C38, C108) and FISNA (C150, C196) were unique to NLRP3 whereas the NBD (C279, C280) and WHD (C514) residues were shared with only a couple other NLRP family members. Thus, the reactive Cys residues in the LRR are highly conserved with other NLRPs while those in the NACHT and PYD domains are not. This suggests that reactive Cys residues in PYD and NACHT may mediate inflammasome responses specific to NLRP3, whereas reactive Cys residues in the LRR may support inflammasome functions common to the entire NLRP family, possibly as a general sensor for cellular redox. The PYD and NACHT of NLRP3 harbour unique sequence motifs that orchestrate specific agonist responses and have critical functions in NLRP3 inflammasome activation [6, 9, 10]. Interestingly, it is these domains and not the LRR that contain the minimally conserved reactive Cys residues. This conclusion is supported by studies that found the LRR to be dispensable for NLRP3 activation [40]. Despite this, other reports have shown that the LRR does support NLRP3 inflammasome activation, so the contribution of LRR to the overall activation of NLRP3 should not be disregarded [7, 9, 41].

It is intriguing that the reactive C150 and C196 residues reside in the FISNA domain since several reports have delineated the novel role played by the FISNA in the activation of NLRP3. Importantly, Tapia-Abellan *et al.* found that potassium deficiency, a common downstream activator of NLRP3 that is associated with ROS production, was sensed by FISNA; furthermore, this region was required to induce the conformational rearrangement of the NACHT to an open, active state [42]. More recent structural data have revealed how the FISNA directly supports the assembly of active NLRP3 oligomers [14]. The FISNA is partially disordered in the inactive NLRP3 structure but becomes fully ordered upon activation, wherein it acts to bridge neighbouring NLRP3 molecules and stabilize the active inflammasome disk. The C150 and C196 residues exist in this flexible and exposed region of the NACHT domain which suggests their oxidative post-translational modification may act as the initiating event that unlocks the inactive NLRP3 cage. Accordingly, previous research identified C150Y as a hyperactive gene variant associated with Cryopyrin-associated Periodic Syndromes (CAPS) [43]. Additional investigation found the C150Y mutant to be resistant to MCC950 (CRID3) inhibition [15]. MCC950 binds multiple subdomains of the NACHT and locks the NLRP3 protein in a catalytically inactive state [44, 45]. Substitution of Tyr would introduce a large aromatic side chain into the FISNA which may induce its conformational transition to the active conformation and subsequently prevent MCC950 ligation. It is tempting to speculate that sulfenylation of C150 may have a similar effect. Interestingly, S740 variants are also well documented in CAPS patients (NCBI ClinVar Database, Variation ID 1694433); this Ser residue was predicted to form a stable hydrogen bond with oxidized C770 in the MD simulation (**Supplemental Figure S4).** However, S740R mutants were previously assessed by Feng *et al*. in an ASC speck formation assay, and they found no significant impacts on nigericin or temperature sensitivity [15].

How reactive Cys residues in the FISNA (and elsewhere in the NACHT) contribute to the molecular rearrangement required for NLRP3 activation is not known. However, there is demonstrable change in the FISNA topology, and hence reactive Cys residue positioning, in the active structure of NLRP3 (8EJ4, NEK7-licenced with ATPγS ligation [14]). While C514 remained similarly positioned in both active and inactive structures, C150, C196, C279 and C280 residues exhibit substantive intramolecular movement with relocation to the opposite side of the NLRP3 structure (**Supplemental Figure S6**). Once the FISNA domain assumes its active confirmation, loop 1 provides direct interaction with the NACHT of a neighbouring NLRP3 subunit to stabilize the active inflammasome disc [14]. Interestingly, the C150 and C514 residues exhibit intermolecular bonding that would help to stabilize the NACHT^A^-NACHT^B^ interface in the active cryo-EM structure. In a “reciprocal handshake”, the C150 sidechain of NACHT^A^ engages with H364 in the NBD of NACHT^B^, and the C514 sidechain in the β-loop of the WHD from NACHT^B^ engages with R157 of loop 1 in the FISNA of NACHT^A^. In addition, the other Cys residues provide intramolecular contacts to stabilize the active FISNA conformation: C279 with Y258 and Y140 (helix 1), C280 with Y136 (helix 1), as well as C196 with K194 and S198 (helix 2).

MD simulation found that Cys oxidization in the LRR disrupted interface interactions that may destabilize the inhibitory cage-like structure of NLRP3 (7PZC), and thereby, could promote inflammasome activation. Given the high degree of conservation for reactive cysteines residing in LRRs of the NLRP family, it is reasonable to speculate that perhaps other NLRPs are also maintained in an inhibited state that is released, at least in part, by oxidation of cysteines. Several other NLRPs that share reactive Cys conservation in the LRR are regulated by cellular ROS (reviewed in [17]). So, it is possible that the LRR domains in these proteins may also function as ROS sensors to induce conformational changes that disrupt autoinhibition. However, further experimentation is required to show if these NLRPs are in fact autoinhibited by the LRR and if oxidation releases this regulatory mechanism.

Although the reactive Cys residues identified in NLRP3 were in positions of impact, it is not yet possible to confirm the direct contributions of these transient modifications to the change in the oligomeric conformation. MD simulation was useful in defining the structural instability generated in the LRR domain following Cys-SOH accumulation. But it would be computationally demanding to attempt modeling of the dynamic process of activation (i.e., from inactive cage to active disc). As a limitation, the model used for MD simulation was a trimer of LRR domains extracted from the inactive decameric complex (7PZC). Positional restrains were applied to simulate the quaternary environment; however, these approximations still represent major sources of potential artifactual bias. Therefore, the findings from the MD simulation should be considered with these caveats in mind. One future approach to assess S-sulfenylation dependent effects on NLRP3 would be to generate point mutants. However, unlike some PTMs such as phosphorylation, there are no accurate mimics for S-sulfenylation that would allow for precise interrogation of each Cys independently. The substitution of acidic Asp/Glu residues, although previously used [46], offers imprecise mimic of sulfenes since it introduces a negative charge state and hydrophilicity similar to sulfinic/sulfonic acids (i.e., non-reversible oxidation) [47]. Substitution of Cys with Ser would replicate the hydroxyl; however, the absence of the sulfur atom notably alters the side chain chemistry relative to a sulfene.

Collectively, this study provides novel understanding of how NLRP3 senses ROS via reactive cysteines and how those oxidative events might impact protein function during inflammasome activation. Numerous reactive cysteines in NLRP3 were identified with *in vitro* and *in situ* labelling approaches. Several reactive Cys were shown to support inflammasome activation in an artificial inflammasome reconstitution. MD simulation suggests LRR oxidation can disrupt LRR-LRR interfaces in the inactive cage. This model is supported by findings that NLRP3-C933A and NLRP3-C941A mutants exhibited reduced IL-1β production in HEK cells expressing inflammasome components. While intriguing observations were made for the location of reactive Cys residues in structures of the NACHT domain, further empirical experimentation is required to validate their mechanistic impact during the activation NLRP3.

## 5. ACKNOWLEDGEMENTS

This work was supported by grants from the Natural Sciences and Engineering Research Council of Canada (NSERC; RGPIN/04379-2019 to JAM, RGPIN 2017-04879 to DCS) and the Canada Foundation for Innovation (CFI; #201818 to JAM, #43143 to DCS). BKZ was supported by a Libin Cardiovascular Institute Doctoral Scholarship and an Alberta Graduate Excellence Scholarship.

## Supplemental Figures

**Supplemental Figure S1.**
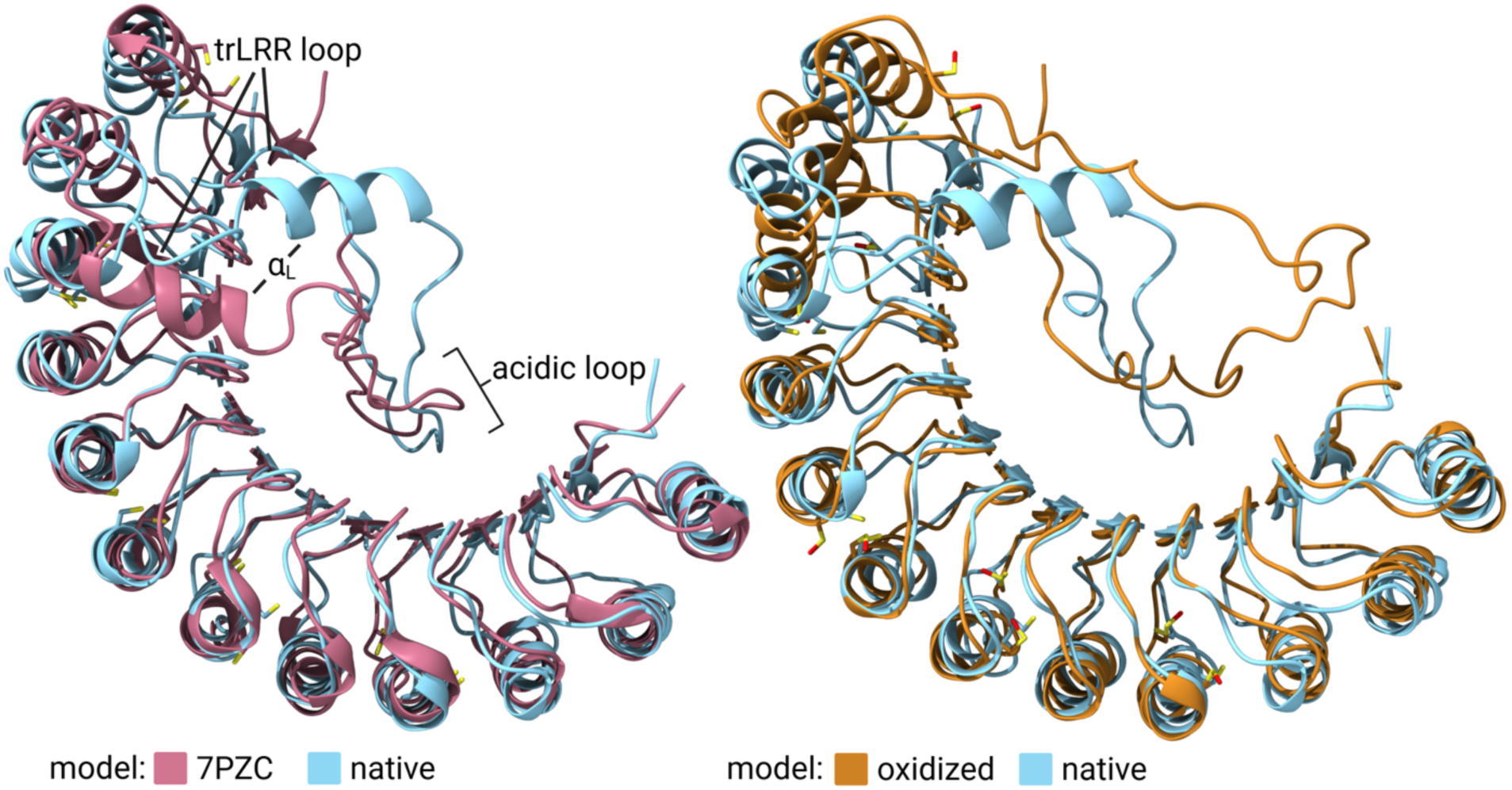
As described in the Materials C Methods, the sidechains of 10 reactive Cys residues in Chain A were converted to Cys-SOH prior to the MD simulation. The final LRR structures are shown at the end of the MD simulation (60 ns). Left: the original LRR structure (Chain A, 7PZC) solved by cryo-EM is overlayed with the final MD simulation model (native) where no alterations to Cys residue sidechains Right: the final structures of the native and oxidized Cys-SOH models are aligned to show conformational changes to the trLRR loop region, including the αL helix and acidic loop.

**Supplemental Figure S2.**
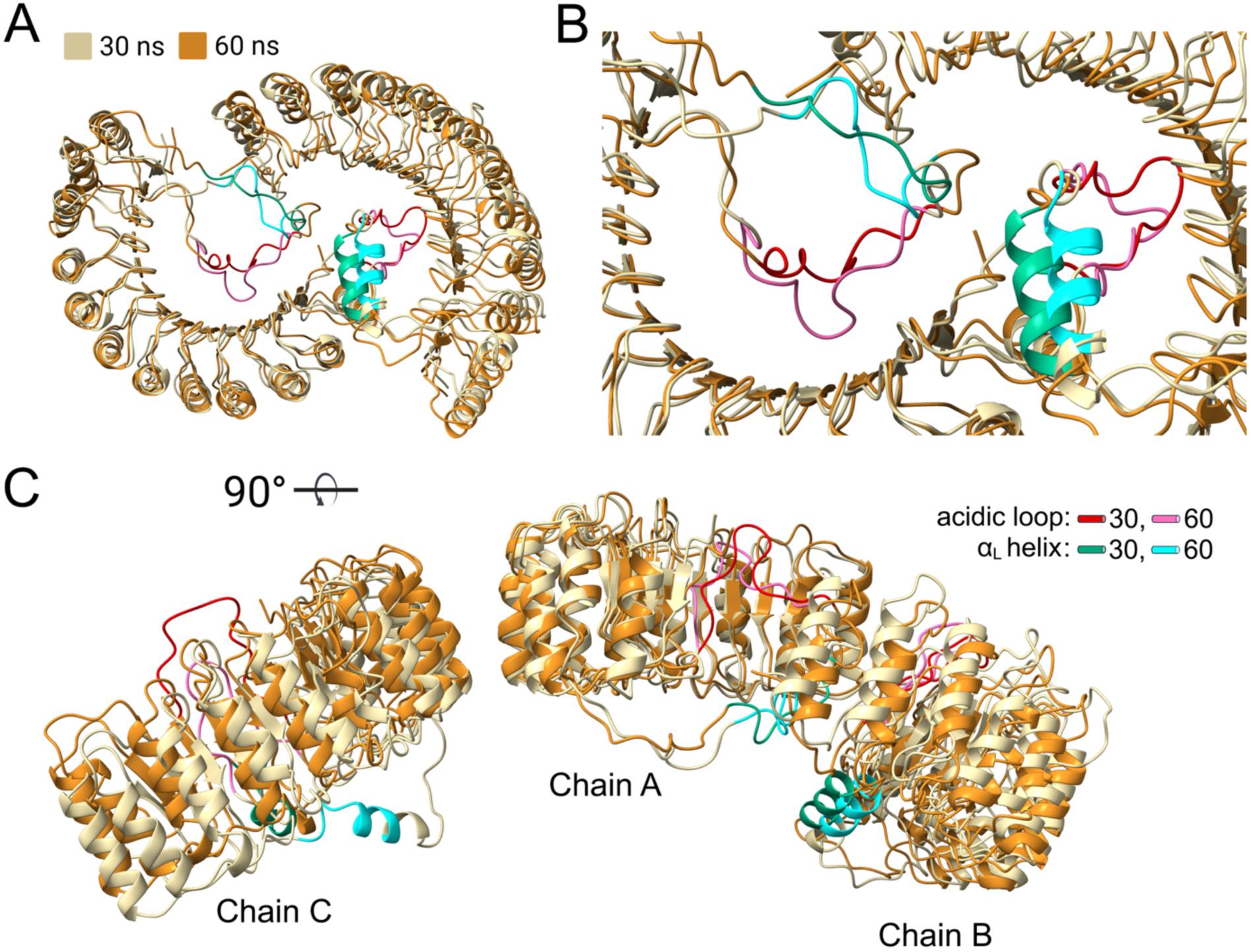
Representative structures of the oxidized Cys-SOH LRR domain model were sampled at 30 and 60 ns during the MD simulation. As described in the Materials C Methods, the sidechains of 10 reactive Cys residues in Chain A were converted to Cys-SOH prior to the MD simulation. Structures were sampled at 30 and 60 ns and then aligned to show conformational changes in: (**A**), the two interacting LRR domains (Chains A and B) and the interface A region within the domain concavities; (**B**), the trLRR loop region, including the αL helix and acidic loop, of interface A; and (**C**), the αL helices and acidic loops of Chains B and C that result from Cys residues oxidation in Chain A.

**Supplemental Figure S3.**
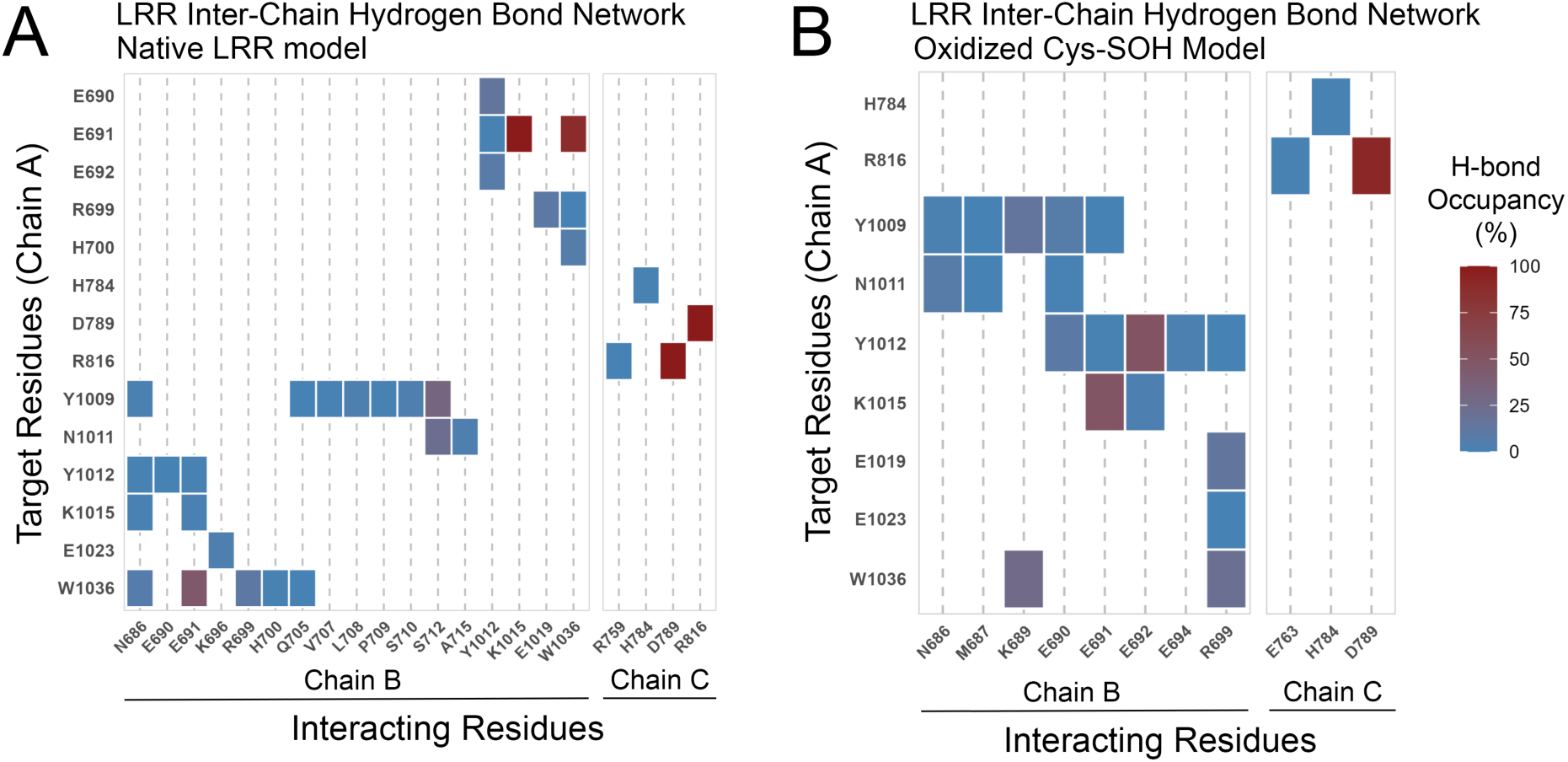
Inter-chain hydrogen bonding (acceptor and donor) for interface residues of native (**A**) and oxidized Cys-SOH (**B**) LRR models. Average hydrogen bond occupancy for the 30 – 60 ns window was calculated in GROMACS.

**Supplemental Figure S4.**
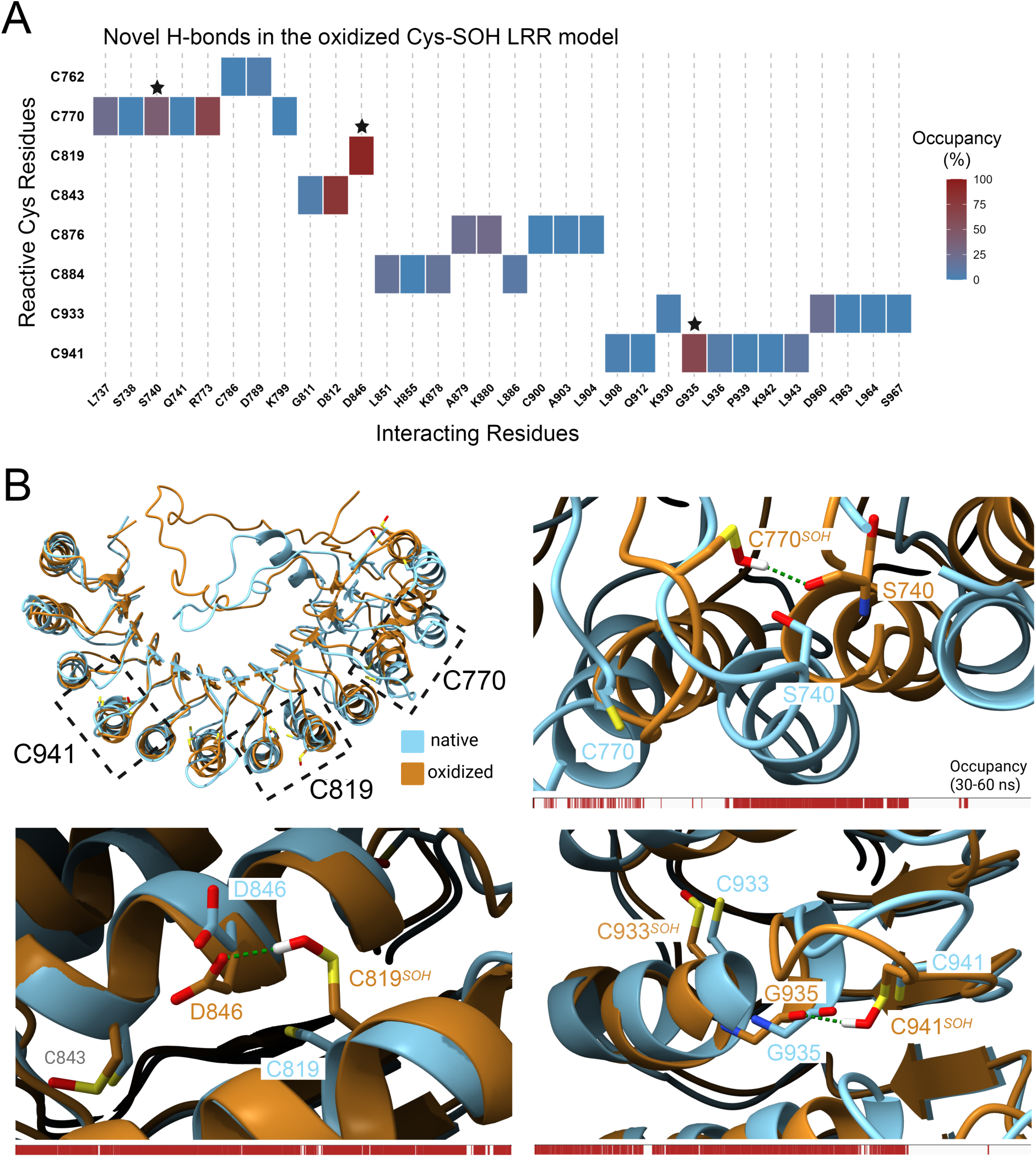
(**A**) Intra-chain hydrogen bonding heatmap of reactive Cys residues (with intra-residue bonds omitted). Black stars (*) indicate hydrogen bonds that are displayed within the structural images. (**B**) Representative images showing novel sulphene hydrogen bonds formed by C770, C819, and C941 in the oxidized LRR model. The red bar below each magnified view shows occupancy for each hydrogen bond over the 30-60 ns time period. All structures were generated using ChimeraX.

**Supplemental Figure S5.**
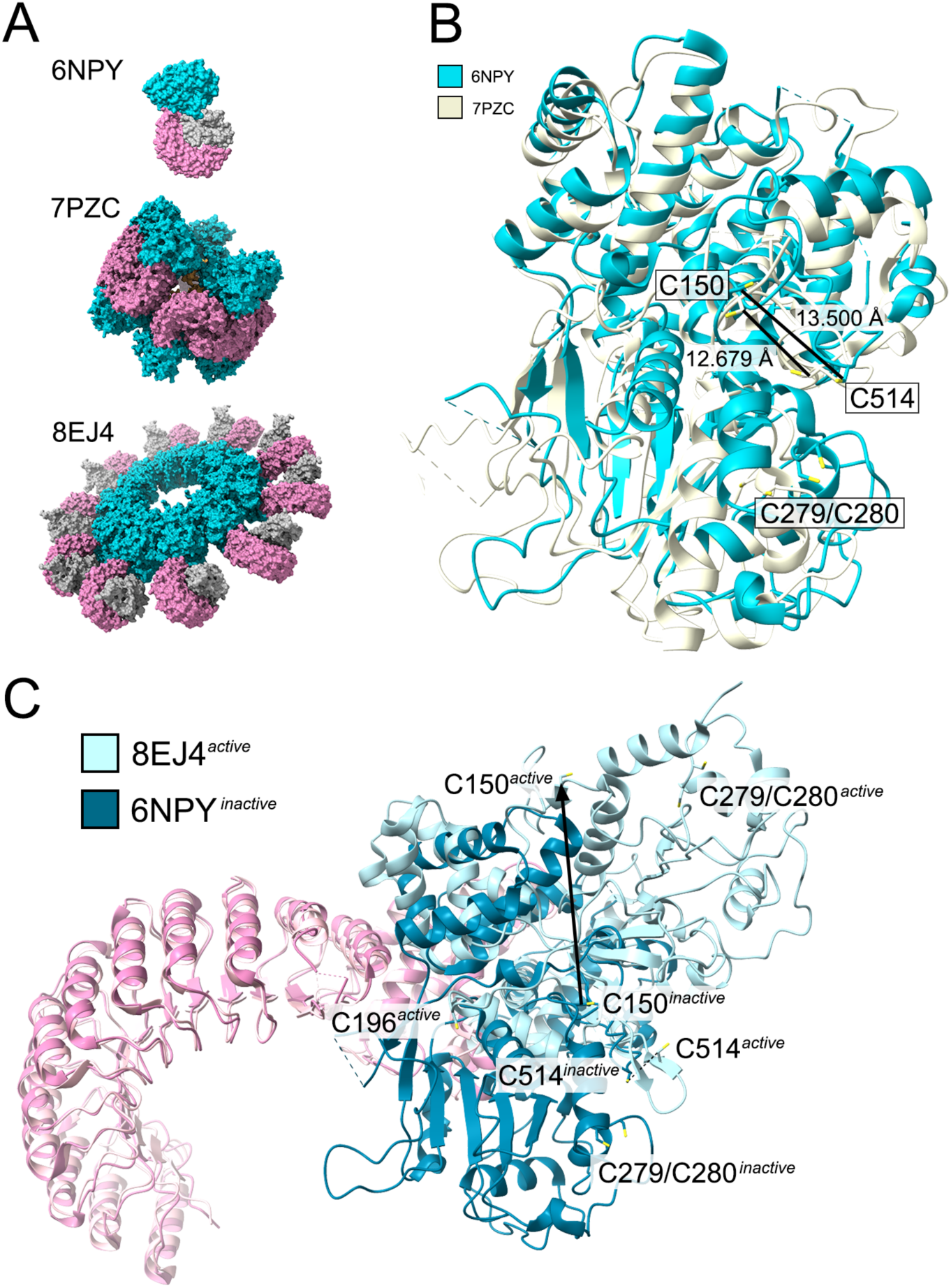
(**A**), structures for inactive (6NPY, 7PZC) and active (8EJ4) NLRP3 were used to highlight the location of the reactive C150, C196, C279, C280 and C514 residues. The NACHT (teal) and LRR (pink) domains are coloured for clarity; NEK7 is shown in gray. (**B**) The NACHT domains from two inactive structures were aligned to show the relative positioning of reactive Cys residues. The NEK7-bound and inactive (ADP-ligated) conformation was obtained from the NLRP3-NEK7 binary complex (6NPY). An inactive (CRID3 ligated) conformation in the absence of NEK7 was obtained from the human NLRP3 cage structure solved by cryo-EM (7PZC). (**C**) The NEK7-bound and active (ATPγS-ligated) conformation was extracted from the human NLRP3 structure solved by cryo-EM in the disc-shaped inflammasome complex (8EJ4) and aligned with the inactive 6NPY structure. All structures were generated using ChimeraX.

**Supplemental Figure S6.**
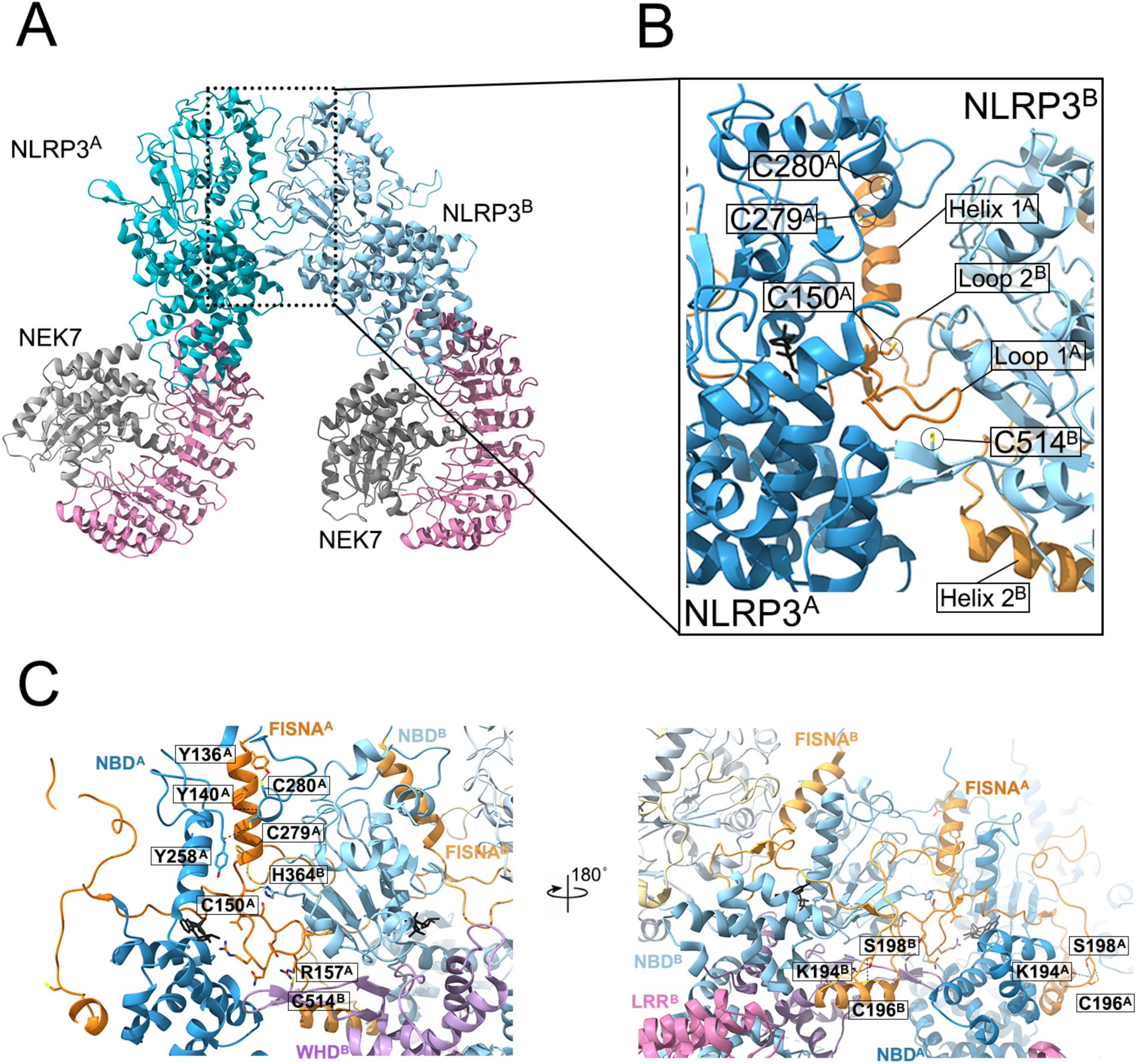
Reactive Cys residues are found at the interaction interface of neighbouring NLRP3 molecules in the active disc. (**A**) The structural representation of two neighbouring NLRP3 molecules was extracted from 8EJ4. (**B**) A close up of reactive Cys residues located in the NLRP3^A^-NLRP3^B^ interface region is provided. The FISNA and NACHT domains are shown in brown and blue, respectively. Important structures of the FISNA domain are labelled. (**C**) Mapped interactions of reactive Cys residues C150, C279, C280 and C514 are shown at the NLRP3^A^-NLRP3^B^ interface. Predicted H-bonds are included along with the ATPγS molecule (black). Mapped interaction of reactive C196 are shown in two neighbouring NLRP3 molecules. All structures were generated using ChimeraX.

